# Motor System Oligodendroglia Atlas Reveals Activation States Associated with Region-Specific Vulnerability in ALS

**DOI:** 10.64898/2025.12.09.693234

**Authors:** Maria Georgopoulou, Frederik Hobin, Jonas Dubin, Pegah Masrori, Geethika Arekatla, Koen Poesen, Sandrine Da Cruz, Ludo Van Den Bosch, Dietmar Rudolf Thal, Alejandro Sifrim, Philip Van Damme

**Author notes:** Senior author (Alejandro Sifrim, Philip Van Damme).

## Abstract

Oligodendrocytes, the second most abundant cell type in the motor cortex and the most abundant in the spinal cord—are increasingly recognized as active contributors to neurodegenerative diseases, including amyotrophic lateral sclerosis (ALS). Yet their diversity and disease relevance in the central nervous system remain incompletely defined. Here, we generate a comprehensive single-nucleus transcriptomic atlas of 176 samples, integrating 2 in-house and 16 publicly available datasets. It encompasses more than *1,000,000* cells (*350,000* oligodendrocytes and oligodendrocyte precursors) from the human motor cortex and spinal cord in non-neurological controls and ALS patients. We identify a previously unrecognized oligodendrocyte subpopulation—marked by *PPEF1*—and uncover regional differences, with distinct OPALIN+ and RBFOX1+ oligodendrocyte proportions between cortex and spinal cord. Importantly, we demonstrate that “disease-associated oligodendrocytes” (DAOs) represent a distinct activation state, rather than a distinct subpopulation. In ALS, spinal cord oligodendrocytes selectively shift toward reactive states, characterized by *JUND*-enriched regulon activity, heightened metabolic demand, increased intercellular signaling, and activation of cell-death pathways. Finally, we identify *LRP2* as a novel transcriptional oligodendrocyte marker and find LRP2 and MBP proteins to be elevated in ALS cerebrospinal fluid (CSF), offering a potential biomarker of oligodendrocyte dysfunction. Our study provides a systems-level framework to interpret oligodendrocyte and OPC changes in ALS, linking cell state shifts to molecular alterations that can guide biomarker and therapeutic strategies.

## INTRODUCTION

Amyotrophic lateral sclerosis (ALS) is a neurodegenerative disorder that primarily affects the motor system, resulting in degeneration of upper motor neurons in the motor cortex and lower motor neurons in the brainstem and spinal cord.^1^ This leads to progressive muscle weakness and wasting, ultimately resulting in respiratory failure.^1^ Median survival from symptom onset is approximately two to five years in the absence of effective therapies.^1,2^ Although more than 40 genes, including *C9orf72*, *SOD1*, *FUS*, and *TARDBP*, have been implicated in ALS, the majority of cases lack an identifiable mutation, and the pathogenic mechanisms remain incompletely understood.^1,3–5^ Neuropathologically, ALS is defined by the cytoplasmic aggregation of TDP-43 (transactive response DNA-binding protein of 43 kDa) in approximately 97% of patients, which is thought to contribute to disease via nuclear loss-of-function and cytoplasmic gain-of-function mechanisms.^6,7^

ALS is recognized as a non-cell-autonomous disease in which glial cells, including astrocytes, microglia, and oligodendrocytes, actively contribute to motor neuron degeneration.^8–13^ Oligodendrocytes are the main myelinating cells in the central nervous system (CNS) and are derived from oligodendrocyte precursor cells (OPCs).^14^ They are the predominant cell type in the spinal cord and the second most abundant in the cerebral cortex, following neurons.^15^ Oligodendrocytes are not only responsible for myelination but also play a key role in metabolic and trophic support of neurons and in immuno-inflammatory responses.^13^ Their dysfunction, including TDP-43 pathology and impaired differentiation from OPCs, has been observed in ALS and precedes motor neuron degeneration in animal models.^9,12,13^ Selective removal of mutant SOD1 from oligodendrocytes substantially delays disease onset in mice, highlighting their causal role in pathology.^9^ Yet, the precise mechanisms and disease-associated states of oligodendrocytes in ALS remain poorly characterized.

Transcriptomic studies in multiple neurological conditions have described “disease-associated” (DA) states in glial cells, including microglia, astrocytes, and more recently, oligodendrocytes.^16,17^ Disease-associated oligodendrocytes (DAOs) are characterized by inflammatory and immune pathway upregulation alongside downregulation of cholesterol biosynthesis, with shared markers including APOD, FTL, and FTH1.^18^ While DAOs have been observed in diseases such as multiple sclerosis, Parkinson’s disease, and Alzheimer’s disease, as well as in trauma^18–26^, they have not been systematically studied in ALS.^27,28^ Moreover, although not classified as DAOs, oligodendrocyte dysfunction has also been noted in other conditions, such as schizophrenia and cancer.^29–32^ However, it remains unclear how diverse oligodendrocyte responses across diseases relate to each other, and whether they follow common organizational principles.

To address this gap, we established a comprehensive single-nucleus RNA-seq (snRNA-seq) atlas of oligodendroglia in human ALS by integrating 18 datasets, both in-house^33^ and publicly available^15,34–47^, covering the motor cortex and spinal cord (cervical, thoracic, and lumbar). This large-scale dataset allowed us to define a unified framework in which heterogeneous disease-associated oligodendrocyte states can be grouped into two activation states, each driven by distinct transcriptional programs. Such a framework not only clarifies oligodendrocyte dysfunction in ALS but also suggests a generalizable approach for studying glial responses across neurodegenerative, demyelinating, traumatic, and psychiatric conditions. Finally, to explore translational potential, we quantified oligodendrocyte-specific proteins in the cerebrospinal fluid (CSF) of ALS patients, assessing their utility as markers of oligodendrocyte loss. Together, these efforts provide both a conceptual framework and a translational entry point for understanding and tracking oligodendrocyte dysfunction in ALS.

## RESULTS

### A Comprehensive Atlas Across Multiple Datasets

To create a motor system oligodendroglia atlas, we conducted single-nucleus RNA sequencing (snRNA-seq) in-house on motor cortex and spinal cord samples. Our dataset comprised 15 samples from both tissues across three conditions—five control, five C9orf72 ALS (C9ALS), and five sporadic ALS (sALS)—yielding a total of 30 samples.^33^ We also integrated our dataset with 16 publicly available motor cortex and spinal cord datasets, resulting in a merged dataset of 176 samples (100 controls, 33 C9ALS, 41 sALS, 1 fALS [familial ALS with unknown mutation], and 1 sALS-FTD [sporadic ALS with frontotemporal dementia comorbidity]). For the motor cortex, we used all nine publicly available snRNA-seq datasets, four of which contained both control and ALS samples, and the remaining five only control.^15, 34–41^ The final dataset, post-integration, consisted of *712,970 cells*, of which *163,906* were oligodendrocytes and *28,947* OPCs (*Figure 1A-1B and S1A-S1B*). For the spinal cord, we used seven published datasets from adult human spinal cord, including one ALS dataset.^15, 42–47^ Post-integration, the spinal cord dataset consisted of *297,688 cells*, of which *146,150* were oligodendrocytes and *15,926* OPCs. Overall, we constructed a comprehensive transcriptomic atlas of the human motor central nervous system, comprising a total of *1,010,658* cells and 13,746 common features, including *310,056* oligodendrocytes and *44,873* OPCs (*Figure 1A-1B and S1A-S1B; Table S1*).

**Figure 1.**
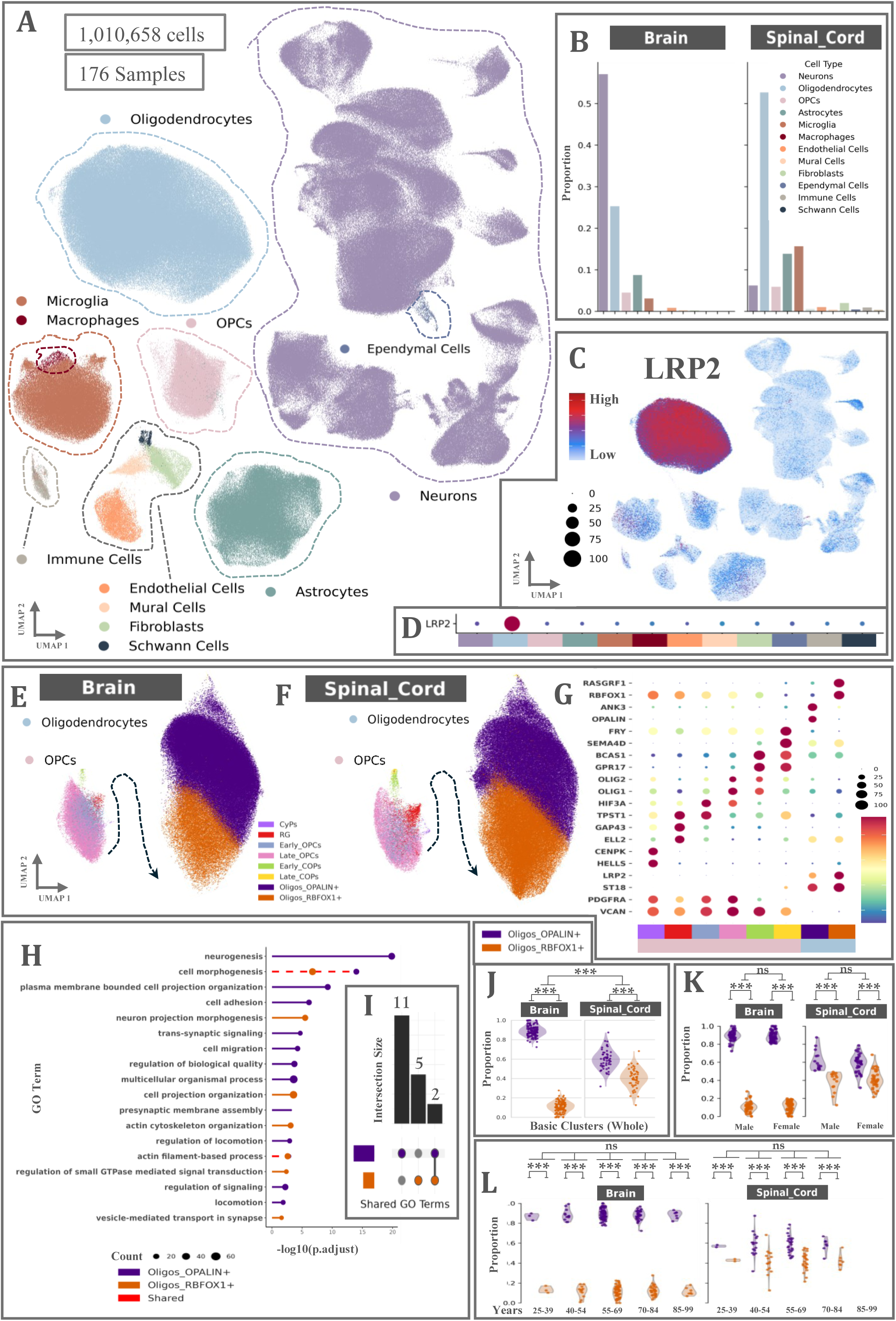
Transcriptomic Atlas of the Motor Cortex and the Spinal Cord Across all Integrated Datasets. **(A)** UMAP the merged motor cortex and spinal cord dataset. **(B)** Proportions of cells of the merged motor cortex and spinal cord dataset. **(C,D)** FeaturePlot **(C)** and DotPlot **(D)** of the expression of LRP2 in the merged motor cortex and spinal cord dataset. **(E-G)** UMAP **(E,F)** and DotPlot **(G)** of the basic clusters and the expression of their marker genes found in OPCs and oligodendrocytes in the motor cortex and the spinal cord. **(H)** GO terms enriched in Oligos_OPALIN+ (purple) and Oligos_RBFOX1+ (orange). The red dashed line represents the common terms. **(I)** Upset plot of the GO terms in the basic clusters. **(J)** Proportions of Oligos_OPALIN+ and Oligos_RBFOX1+ clusters in the merged control motor cortex and spinal cord dataset. Each point is a Sample_ID. **(K)** Proportions of Oligos_OPALIN+ and Oligos_RBFOX1+ clusters in the merged control motor cortex and spinal cord dataset split by sex. Each point is a Sample_ID. **(L)** Proportions of Oligos_OPALIN+ and Oligos_RBFOX1+ clusters in the merged control motor cortex and spinal cord dataset split by age group. Each point is a Sample_ID. *** p-value < 0.001 ns: not-significant See also **Figure S1**.

Quality control metrics—such as the percentage of mitochondrial reads, total counts, and the number of unique features—demonstrated consistent results across all datasets. Across all datasets, approximately 91–94% of cells per sample passed quality control, yielding *712,970* high-quality cells for downstream analyses. The proportion of retained cells was comparable across control and ALS samples, indicating no systematic bias was introduced during filtering. Evaluation with scIB metrics on the UMAP space showed consistently low rejection rates among clusters, indicating robust local batch mixing. These results support that harmony integration successfully minimized batch effects. (*Table S2*).^48^

Based on gene expression, we first classified all cells into 12 primary “Cell Types” based on established marker genes (*Figure 1A and S1C-S1D*). Within the oligodendrocyte population, <u>Low-Density Lipoprotein</u> <u>Receptor-Related Protein 2</u> (LRP2, also known as Megalin) emerged as a candidate lineage marker, whose expression increases progressively with maturation (*Figure 1C-1D*). Previous studies have highlighted that megalin has an important role in OPC and oligodendrocyte maturation.^49–51^ Here, we propose LRP2 as a specific marker for oligodendrocytes, suitable for lineage annotation and quantification (*Figure 1C-1D*).

With more granular subclustering, we captured distinct basic cell populations, called “Basic Clusters”, representing cell states within each primary type defined by known markers. While we performed detailed subclustering and activation state annotation for oligodendrocytes and OPCs, annotations of other glial cell types for reactive or homeostatic states remained exploratory. Detailed subclustering and functional analysis of these other cell types were beyond the scope of this study and thus are not included in the main analyses.

Focusing on oligodendroglial subpopulations, we identified 8 distinct basic clusters (*Figure 1E-1G*): 6 immature populations, 2 mature oligodendrocyte populations, *OPALIN*+ (Oligos_OPALIN+) and *RBFOX1*+ (Oligos_RBFOX1+) (*Figure 1E-1G*). GO analysis of the Oligos_OPALIN+ and Oligos_RBFOX1+ populations show that they share many GO terms. Among the terms that are unique to each population, Oligos_RBFOX1+ is enriched in processes related to regulation of actin cytoskeleton organization, small GTPase-mediated signal transduction, and vesicle-mediated transport at the synapse (*Figure 1H-1I*). Oligos_OPALIN+ cells are enriched for GO terms associated with glial and neuronal development and differentiation, cell adhesion and migration (*Figure 1H-1I*).

Interestingly, the distribution of oligodendrocyte subpopulations was region-dependent (lmm; accounting for donor level variability, p<0.001). In the motor cortex, Oligos_OPALIN+ comprised more than 80% of the total oligodendrocyte population, with the remaining ∼20% being Oligos_RBFOX1+ (*Figure 1J*). In contrast, in the spinal cord, these proportions were nearly equal, suggesting a difference in the abundance of actively myelinating oligodendrocytes between the motor cortex and spinal cord, which we confirmed across individual datasets (*Figure 1J*). Although it has been previously suggested that Oligos_OPALIN+ accounted for >80% and Oligos_RBFOX1+ for <20% in the spinal cord^15^, the large number of control samples in our study (100 controls) allowed us to refine these estimates, suggesting that the two subpopulations are instead present in roughly equal proportions. We next examined potential age (*Figure 1K*) and sex (*Figure 1L*) effects on basic cluster proportions. No significant associations were detected (*Figure 1K-1L*), suggesting that regional differences within the CNS represent the predominant determinant of the abundance of these oligodendrocyte populations.

We further subclustered oligodendrocytes and OPCs to define distinct cell states based on marker expression. Leveraging the scale of the dataset, we refined and extended previous single-dataset subclustering efforts^19,52^, providing a unified and comprehensive atlas of oligodendroglial subclusters. We identified 14 subpopulations, hereafter referred to as “Subclusters” (7 OPC subclusters and 7 oligodendrocyte subclusters), in the motor cortex and spinal cord (*Figure 2A-2G; Table S4-S5*). These subclusters represent a continuum of transition states from early to late maturation oligodendroglial populations. Manual inspection using subsetting, differential marker analysis, and post-hoc annotation confirmed that these states captured the data’s complexity. After correction for batch effects and donor variability, all subclusters remained robustly represented across samples, datasets, and conditions.

**Figure 2.**
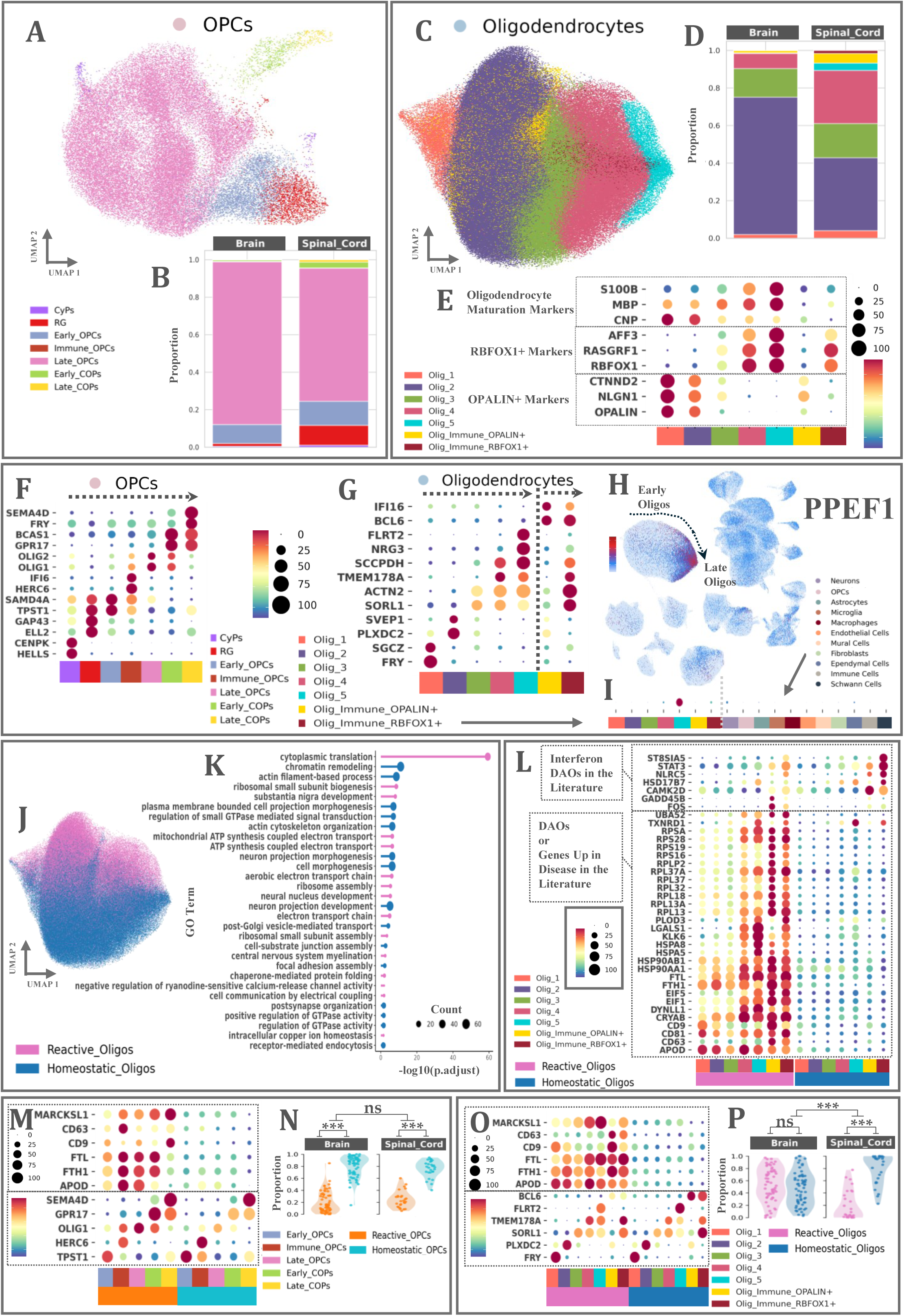
Oligodendrocyte and OPCs Subclusters and Activation States. **(A)** UMAP of the subclusters in the OPC population. **(B)** Proportions of the subclusters in the control OPC population. **(C)** UMAP of the subclusters in the oligodendrocyte population. **(D)** Proportions of the subclusters in the control oligodendrocyte population. **(E)** DotPlot of the expression of known oligodendrocyte RNA/protein maturation markers and Oligos_OPALIN+ / Oligos_RBFOX1+ specific markers across oligodendrocyte subclusters. **(F and G)** DotPlot of the top marker genes of each OPC **(F)** and oligodendrocyte **(G)** subcluster. **(H and I)** FeaturePlot **(H)** and DotPlot **(I)** of the expression of the Olig_5 specific marker PPEF1 in the merged motor cortex and spinal cord dataset. In the plot **(I)** the groups are clustered by cell population. The oligodendrocyte population is further grouped by subcluster. **(J)** UMAP of the activation states in the oligodendrocyte population in the of the merged motor cortex and spinal cord dataset. **(K)** GO terms enriched in reactive oligodendrocytes (pink) and homeostatic oligodendrocytes (blue). The red dashed line represents the common terms. **(L)** DotPlot of the gene expression in each subcluster spitted by activation state. The features are selected reactive or disease-associated or interferon disease-associated markers from the literature. **(M and O)** DotPlot of the gene marker expression in each subcluster spitted by activation state in the OPCs **(M)** and the oligodendrocytes **(O)**. In the top part of the DotPlots are the main activation state marker genes (*APOD*, *FTL*, *FTH1*, *CD9*, *CD63*, *MARCKSL1*). In the bottom part are the subcluster specific genes. **(N and P)** Proportions of reactive states and homeostatic states in the control OPCs **(M)** and oligodendrocytes **(P)**. Each point is a Sample_ID. Abbreviations in this figure include: OPCs; Oligodendrocyte Progenitor Cells. *** p-value < 0.001 ns: not-significant

We consistently observed a previously unreported subpopulation, Olig_5, which uniquely expresses *PPEF1* (*Figure 2H-2I*). The Olig_5 subcluster was detectable but scarce in the motor cortex compared to the spinal cord, due to the lower abundance of Oligos_RBFOX1+ in the cortex. This subpopulation is unlikely to be a technical artifact, as it was conserved across all datasets, shows normal QC metrics and does not express markers of other cell types.

### Reactive and Homeostatic Activation States Across Health and ALS

Our unbiased clustering did not reveal a DAO-specific cluster. DAOs have been implicated in various conditions, from neurodegenerative diseases to trauma. Canonical markers of this cell state include *FTL*, *FTH1*, *APOD*, *CD9*, *CD63*, *CRYAB*, *NKX6-2*, and *MARCKSL1*.^18–26^ We annotated cells using these markers via Leiden clustering with post hoc annotation. Additionally, we applied a priori selection of DA marker genes followed by Gaussian Mixture Model clustering, yielding strong agreement between methods.^53^ These analyses indicate that DAO-like cells form a continuous transcriptional state rather than a discrete population (*Figure 2J; Table S6*).

We selected these markers for annotation for two main reasons. First, they are consistently reported in the literature as upregulated in disease-associated states. Second, they showed strong loading on principal component 2 (PC2) within the oligodendrocyte population, whereas PCs 1 primarily captured maturation differences between Oligos_OPALIN+ and Oligos_RBFOX1+. This indicates that DAO markers define an orthogonal axis of variation, independent of maturation (*Figure 2J*).

Importantly, these continuous states were present in both control and ALS samples, indicating they are not exclusively disease-associated but reflect a broader physiological phenomenon. Although they are more prevalent in disease, the term “disease-associated” remains useful but incomplete. To more accurately capture this biology, we classified cells into two activation states. The first group, termed “Reactive”, showed upregulation of DAO-associated markers, whereas the second, termed “Homeostatic”, displayed lower expression of these markers, consistent with a baseline condition. GO analysis (presented below) reinforced these functional assignments. This reframing highlights that so-called DAOs are better understood as general activation states—present in health but usually preferentially enriched in disease.

These populations exhibited distinct distribution and polarization in low-dimensional UMAP projections (*Figure 2J*). Reactive and homeostatic cells span across the maturation continuum, from early Oligos_OPALIN+ to later Oligos_RBFOX1+ stages. Thus, activation is not a separate lineage but a dynamic overlay on the oligodendrocyte maturation trajectory.

To compare the two distinct activation states, we performed differential expression analysis (DEA).^54^ DEA confirmed known DA signatures in both oligodendrocytes and OPCs. In reactive oligodendrocytes, key upregulated pathways included myelination and translation, while homeostatic oligodendrocytes showed enrichment in signaling, synapse formation, and cell development (*Figure 2K*). Only reactive states exhibited upregulation of cell death pathways, consistent with increased susceptibility under both healthy and disease conditions, whereas homeostatic states appeared to play more supportive roles (*Figure 2K*). Similarly, reactive OPCs were enriched in translation, protein processing, and apoptosis pathways, whereas homeostatic OPCs displayed pathway profiles similar to those of homeostatic oligodendrocytes. Together, these findings reveal that reactive states capture stressed, potentially vulnerable oligodendrocytes, while homeostatic states represent supportive baseline functions.

For annotating the activation state, we initially used previously mentioned DAO markers. Additionally, several other genes previously reported^18–26^ as upregulated in DAOs were also consistently enriched across reactive states, supporting the robustness of our clustering (*Figure 2L*). Using this marker framework, we recovered >95% of human DAO-associated genes previously reported. In our classification, genes upregulated in disease mapped to reactive states, those downregulated mapped to homeostatic states, and interferon-responsive DAOs corresponded to either reactive or homeostatic Olig_Immune (*Figure 2L*). This validation demonstrates that our framework captures nearly all known DAO biology while extending it into a more generalizable activation model.

We identified both reactive and homeostatic states within each oligodendrocyte and OPC subcluster, which we annotated as “Subcluster Activation States”. Reactive states expressed the same subcluster-specific markers as their homeostatic counterparts, but additionally showed upregulation of canonical DAO markers (*Figure 2M and 2O*). We did not assign activation states to CyP and RG subclusters, as their disease associations are not well characterized and their low cell numbers raised the risk of technical artifacts. This approach allowed us to dissect activation while preserving developmental context, revealing that reactivity is a feature within, rather than outside of, oligodendrocyte subtypes.

Significant proportional differences (*Figure 2N and 2P*) in oligodendrocyte activation states (*Figure 2P*) were observed between the motor cortex and the spinal cord. However, OPCs did not show such differences (*Figure 2N*). In the control motor cortex, there was a marked proportional increase in reactive oligodendrocytes compared to other conditions, accompanied by substantial variability in the proportions of reactive and homeostatic states (*Figure 2P*). This variability was not observed in cortical OPCs or in either oligodendrocytes or OPCs of the spinal cord. These findings likely reflect the higher plasticity and demand for adaptive remodeling in the motor cortex, as opposed to the more stable environment of the spinal cord, where oligodendrocyte reactivity appears less pronounced.

### Compositional Variations in Oligodendrocyte and OPC Populations in ALS

Building on previous findings from model organisms and immunohistochemistry in humans, we were interested in determining whether ALS alters the abundance of oligodendrocyte and OPC populations^9–10^ (*Figure 3A-3B*). CSF MBP levels (control: n = 17; sALS: n = 10; C9ALS: n = 8) were significantly increased in both ALS groups relative to controls, while LRP2 (control: n = 18; sALS: n = 18; C9ALS: n = 8) was significantly elevated in sALS (*Figure 3A*; *Table S3*). CNP (control: n = 17; sALS: n = 10; C9ALS: n = 10) showed a trend toward higher levels in the disease groups, though this did not reach statistical significance at current sample sizes (*Figure 3A*). Because these proteins are primarily intracellular^49,55,56^, their presence in CSF likely reflects oligodendrocyte injury or turnover, rather than active secretion. Although MBP, LRP2, and CNP are not entirely specific markers of primary oligodendrocyte loss—since their changes may also result from secondary effects of axonal degeneration— the consistent elevations in ALS CSF suggest that oligodendrocyte perturbation is a component of ALS pathology. Supporting this, in the sALS group, CSF MBP and LRP2 positively correlated with neurofilament light chain (NfL) levels (sALS: n = 25; C9ALS: n = 9), and MBP also correlated with phosphorylated neurofilament heavy chain (pNfH) (sALS: n = 25; C9ALS: n = 9), established markers of neuronal/axonal injury (*Figure 3B*).^57,58^

**Figure 3.**
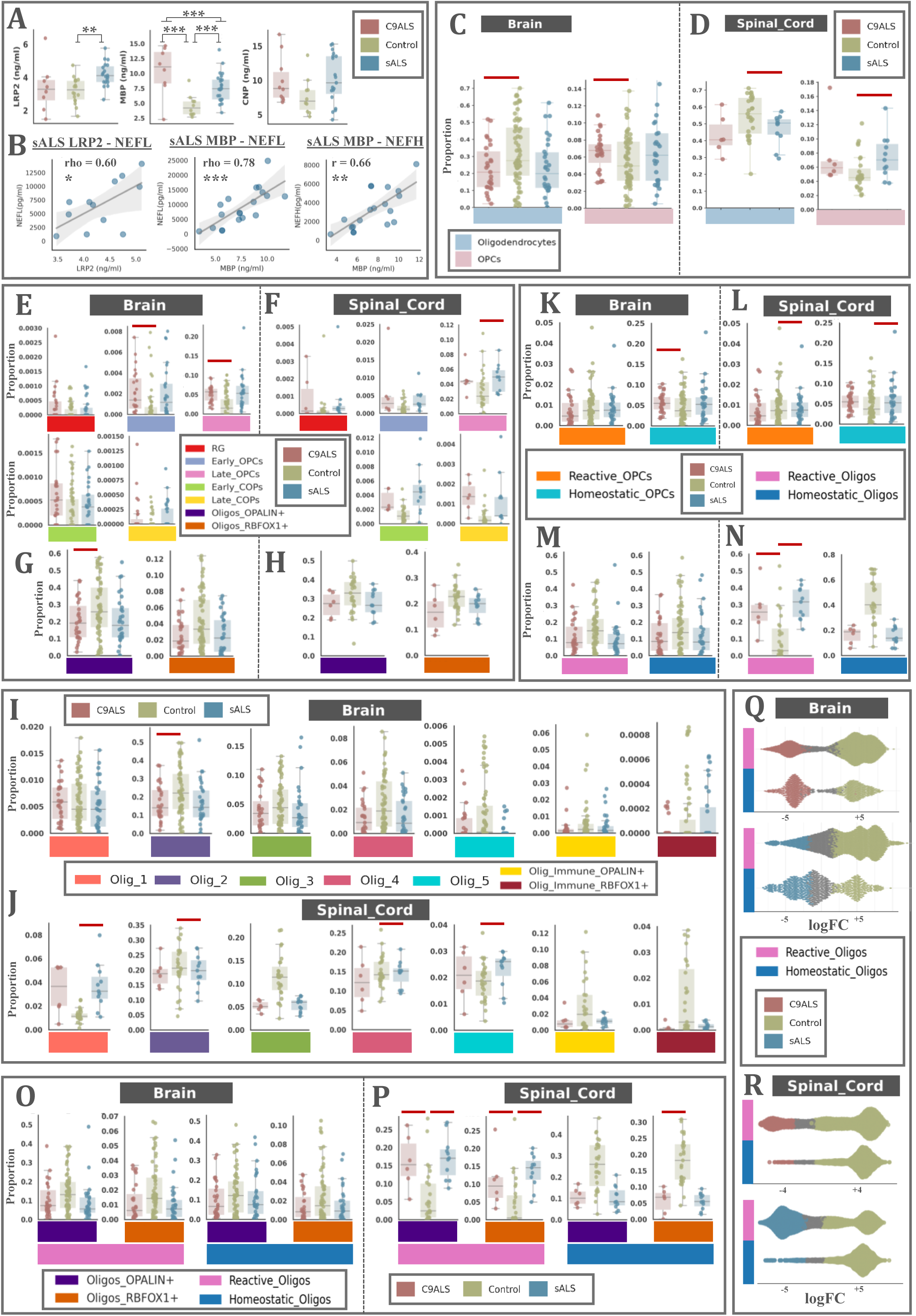
Proportional Differences Between Control and ALS. **(A)** Differences in the levels of LRP2, CNP and MBP in the control, C9ALS and sALS CSF. **(B)** Spearman correlations of the levels of LRP2 and MBP with the levels of NEFL (Nfl) and NEFH (pNEFH) in sALS CSF. **(C and D)** Proportions of OPCs and oligodendrocytes in control, C9ALS and sALS in the motor cortex **(C)** and the spinal cord **(D)**. **(E and F)** Proportions of basic OPC clusters in control, C9ALS and sALS in the motor cortex **(E)** and the spinal cord **(F)**. **(G and H)** Proportions of basic oligodendrocyte clusters in control, C9ALS and sALS in the motor cortex **(G)** and the spinal cord **(H)**. **(I and J)** Proportions of oligodendrocyte subclusters in control, C9ALS and sALS in the motor cortex **(I)** and the spinal cord **(J)**. **(K and L)** Proportions of OPCs activation states in control, C9ALS and sALS in the motor cortex **(K)** and the spinal cord **(L)**. **(M and N)** Proportions of oligodendrocyte activation states in control, C9ALS and sALS in the motor cortex **(M)** and the spinal cord **(N)**. **(O and P)** Proportions of oligodendrocyte basic clusters grouped by activation state in control, C9ALS and sALS in the motor cortex **(O)** and the spinal cord **(P)**. **(Q and R)** Bee swarm plots of oligodendrocyte activation state neighborhoods in control, C9ALS and sALS in the motor cortex **(Q)** and the spinal cord **(R)**. Each dot is one neighborhood. With grey color are shown the not significant ones. Abbreviations in this figure include: OPCs; Oligodendrocyte Progenitor Cells. Each point is a Sample_ID. Proportions were calculated in the whole dataset. Subsetting was applying only for visualization. **(A)** Kruskal–Wallis and Mann–Whitney tests with FDR correction; * p-value < 0.05, ** p-value < 0.01, *** p-value < 0.001 **(C - P)** Credible (scCODA) results, with FDR < 0.05 are depicted as red colored bars.

We next investigated whether ALS affects the relative abundance of total oligodendrocyte and OPC populations, and at the levels of basic clusters, subclusters, and activation states, acknowledging the limitations of single-nucleus sequencing for compositional inference (*Figure 3C-3R*). Disease-specific proportional changes were assessed using multiple Differential Abundance (DA) methods, both cluster-based and cluster-free.^59–63^

For simplicity, the main text presents results from scCODA, which are representative of all methods; all significant findings were corroborated by at least one additional approach. Datasets experimentally enriched for specific cell populations (i.e., samples in which one or more cell types were selectively increased through tissue dissection or nuclear marker–based FACS sorting; labeled as ‘Enrichment == Yes’) were excluded from compositional analyses. This exclusion did not substantially alter overall cell numbers or relative proportions.

In the motor cortex and spinal cord of C9ALS and sALS patients, the relative number of total oligodendrocytes appeared reduced in almost all cases, although this decrease did not always reach statistical significance at current sample sizes (*Figure 3C-3D*). The decrease in oligodendrocytes was accompanied by increased OPC proportions (*Figure 3C-3D*). Specifically, the oligodendrocyte and OPCs proportions were significantly altered in C9ALS motor cortex and sALS spinal cord (*Figure 3C-3D*). Although not all observed changes were statistically robust, the patterns were compatible with a model in which oligodendrocyte loss in ALS is accompanied by compensatory OPC expansion.^9^

This reduction in oligodendrocyte abundance, accompanied by an increase in OPCs, is also evident at the basic cluster level, characterized by a significant expansion of the Early and Late_OPCs populations and a corresponding decrease in the Oligos_OPALIN+ subset relative to controls in the brain (*Figure 3E-3H*).

At the subcluster level in the spinal cord, there was a significant increase in Olig_1 oligodendrocytes in the sALS group—representing early post-OPC stages—along with a reduction in more mature oligodendrocyte populations (*Figure 3I-3J*). This shift supports the hypothesis that OPCs are proliferating in an attempt to compensate for oligodendrocyte loss. Along the OPC-to-oligodendrocyte differentiation trajectory, this trend persists, with an increased abundance of early (Olig_1) oligodendrocyte populations and a decline in more mature populations. Notably, the newly identified Olig_5 population appears resistant to degeneration, as its abundance remains stable compared to the reductions observed in other late-stage oligodendrocytes (*Figure 3J*).

Activation states exhibited region-specific patterns, with different trends in cortex vs. spinal cord (*Figure 3K-3N*). In the motor cortex, there was a modest, but not statistically significant, reduction of both activation states in the disease (*Figure 3K* and *3M*). In contrast, in the spinal cord, oligodendrocytes displayed a significant increase in reactive states, accompanied by a corresponding decrease in homeostatic populations (*Figure 3L* and *3N*).

When activation states were considered within basic clusters, in the spinal cord reactive Oligos_RBFOX1+ and Oligos_OPALIN+ were enriched, whereas their homeostatic counterparts were reduced (*Figure 3O-3P*). These patterns suggest that ALS preferentially affects reactive populations in the spinal cord, while changes in the motor cortex are subtler and variable across datasets. The reduction of both reactive and homeostatic populations in the motor cortex aligned with previous findings describing a decrease in myelination markers such as *CNP*, *MBP*, *SOX8*, *OLIG1* and *OLIG2* in the integrated datasets.^37,38^ These genes, typically enriched in the reactive states, were found decreased in the cortex.

Cluster-free DA analysis, which identifies local regions of transcriptional space enriched for disease, supported our previous observations (*Figure 3Q-3R*). In the spinal cord, ALS-enriched neighborhoods were primarily located within reactive states of oligodendrocytes, whereas no significant differences were detected in the motor cortex (*Figure 3Q-3R*). In both C9ALS and sALS, significant neighborhoods were restricted to reactive states, whereas in controls they spanned both reactive and homeostatic states (*Figure 3Q-3R*).

### Region- and Condition-Specific Transcriptional Oligodendroglial Response to ALS

To identify transcriptional changes associated with ALS, we analyzed both motor cortex and spinal cord tissues using pseudobulk and single-nucleus differential expression analysis.^54,65–67^ We examined both global disease effects (ALS vs. control) and tissue-specific transcriptional responses (motor cortex vs. spinal cord changes). In our dataset, pseudobulk analyses accounted for dataset and sex. These analyses provided a better fit because several cell populations exhibited differential abundance. In contrast, single-nucleus–level tests tended to identify genes as differentially expressed primarily due to changes in cell-type composition rather than true per-cell transcriptional differences. Therefore, the differential expression results shown in the figures are derived from the DESeq2 pseudobulk analysis, with findings cross-validated using Wilcoxon rank-sum test.

#### Motor Cortex vs. Spinal Cord

First, we observed that the motor cortex exhibited significantly fewer differentially expressed genes (DEGs) between ALS and controls compared to the spinal cord, where the number of DEGs was substantially higher (*Figure 4A* and *S2A*). Second, our analysis revealed a markedly smaller number of DEGs in sALS compared to C9ALS, when both were compared with control motor cortex (*Figure 4A* and *S2A*). These results are consistent with earlier protein-level histopathological data, with more pronounced changes in C9ALS compared to sALS cortex.^68^ Finally, our findings in the spinal cord further supported the DA analysis and showed a disease-associated shift toward reactive states, as most of the DEGs could be attributed to this shift (*Figure 4B-4I*).

**Figure 4.**
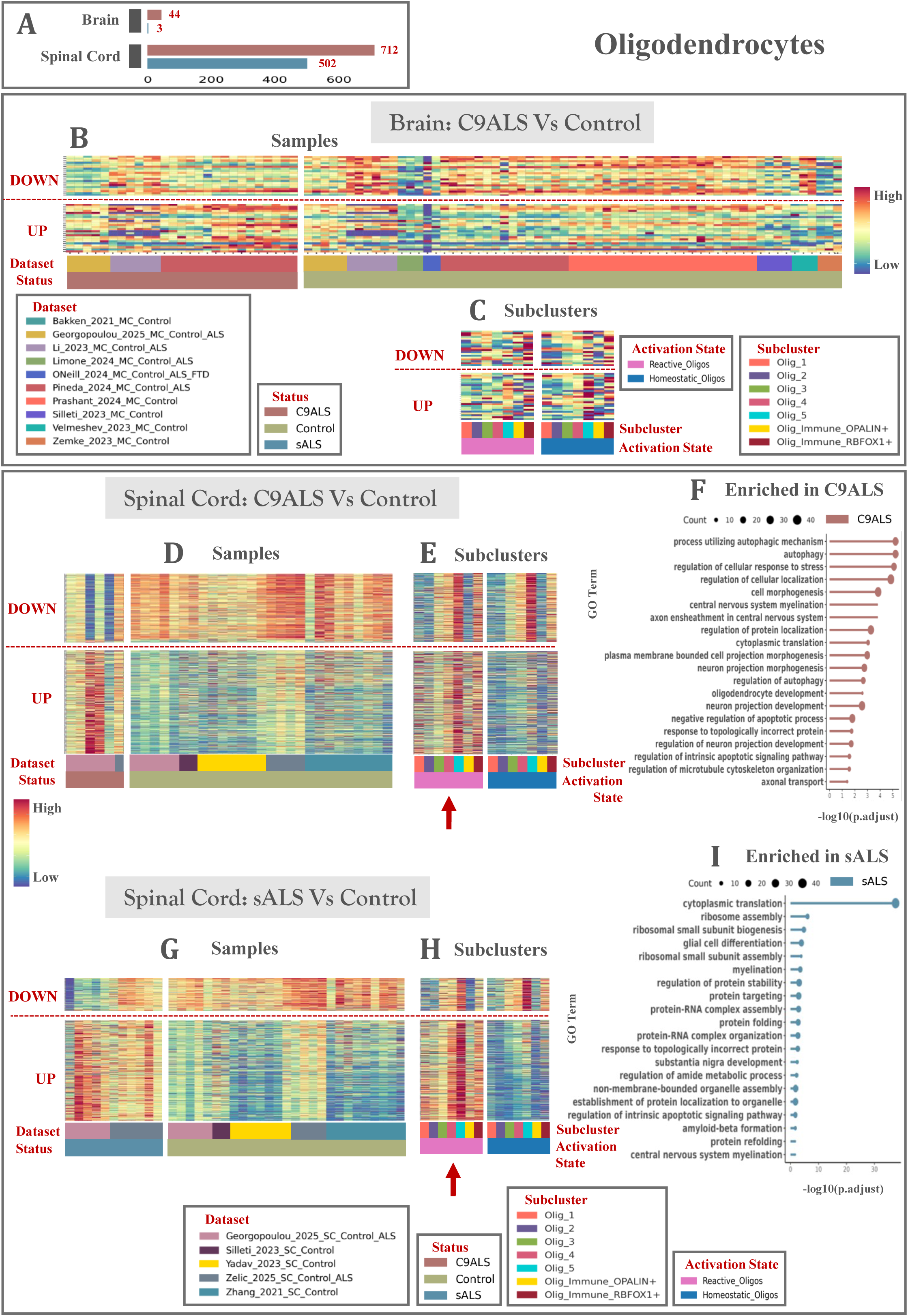
Oligodendroglial Transcriptional Responses to ALS Are Region- and Condition-Dependent. **(A)** Barplot of the amount of significant DEGs found in the motor cortex and the spinal cord with pseudobulk DESeq2 analysis and p.adj < 0.05. **(B)** MatrixPlot of the top up- and down-regulated transcripts in C9ALS vs. control in the motor cortex. Each column is a sample grouped by dataset and status. **(C)** MatrixPlot of the top up- and down-regulated transcripts in C9ALS vs. control in the motor cortex. Each column is a subcluster grouped by activation state. **(D)** MatrixPlot of the top up- and down-regulated transcripts in C9ALS vs. control in the spinal cord. Each column is a sample grouped by dataset and status. **(E)** MatrixPlot of the top up- and down-regulated transcripts in C9ALS vs. control in the spinal cord. Each column is a subcluster grouped by activation state. **(F)** Lollipop Plots of enriched and suppressed GO terms in C9ALS vs. control spinal cord. **(G)** MatrixPlot of the top up- and down-regulated transcripts in sALS vs. control in the spinal cord. Each column is a sample grouped by dataset and status. **(H)** MatrixPlot of the top up- and down-regulated transcripts in sALS vs. control in the spinal cord. Each column is a subcluster grouped by activation state. **(I)** Lollipop Plots of enriched and suppressed GO terms in sALS vs. control spinal cord. Abbreviations in this figure include: p.adj; p adjusted value, DEGs; differentially expressed genes. See also **Figure S2**.

#### Oligodendrocytes sALS vs. control and C9ALS vs. control

In the motor cortex, transcriptional changes were modest. Only a few genes were significantly upregulated (p.adj < 0.05), and this region showed high inter-dataset variability (*Figure 4B–4C*; *Table S7* and *S11*). sALS transcriptomes were largely indistinguishable from controls (*Table S7*). Conversely, in the spinal cord, most DEGs were observed in reactive oligodendrocyte states (*Figure 4D-4E* and *4G-4H*; *Table S9-S10*). Upregulated genes in ALS were enriched in reactive states, while downregulated genes were mainly associated with homeostatic states (*Figure 4E* and *4H*). This indicates a global shift toward a reactive activation state, supporting the notion that oligodendrocytes transition toward a stress-associated phenotype in ALS.

GO term enrichment in the spinal cord revealed upregulation of myelination, cell death pathways, translation and amyloid precursor protein metabolism, consistent with oligodendrocyte stress and potential impairment of myelination (*Figure 4F* and *4I*; *Table S12-S14*). For this analysis, DEGs from pseudobulk DESeq2 with log₂FC > 0.3 and adjusted p.adj < 0.05 were used. Only genes expressed in at least 20% of cells per condition were included to minimize noise and focus on biologically meaningful changes.

#### OPCs sALS vs. control and C9ALS vs. control

Similar to the oligodendrocytes, the motor cortex OPCs, showed minor transcriptional changes in the disease. (*Figure S2B–S2C*; *Table S15*). The disease primarily affects spinal cord OPCs, showing a clear shift in maturation stage (*Figure S2D-S2E* and *S2G-S2H*; *Table S17-S18*). Upregulated genes are mainly associated with the COPs, while downregulated genes are linked to the earlier OPC states (radial-glia like cells, CYPs and OPCs) (*Figure S2E* and *S2H*). This maturation-specific pattern contrasts with mature oligodendrocytes, where such a distinction is absent. Similar to oligodendrocytes, affected OPCs in the spinal cord show enrichment in pathways related to translation, protein metabolism, and myelination (*Figure S2F* and *S2I*; *Table S19-S22*).

#### Oligodendrocytes sALS vs. C9ALS

Both for oligodendrocytes and OPCs comparisons between sALS and C9ALS revealed only a few DEGs in the spinal cord, while in the motor cortex the sALS transcriptome was indistinguishable from controls. Therefore, the contrasts between C9ALS and sALS largely recapitulate those observed between C9ALS and controls in the cortex. For these reasons, we did not pursue additional analyses of these comparisons.

### Dynamic Transition between the Reactive and the Homeostatic States in Health

To investigate the origin of reactive oligodendrocytes and their relationship to homeostatic cells, we applied trajectory inference analyses, constructing high-level maturation trajectories of the oligodendrocyte lineage and arranging cells along pseudotime. We used MARGARET and PAGA to generate an abstract graph of our cell populations, which spanned from early-stage OPC subclusters (radial-glia-like cells, RG) to mature oligodendrocytes (Olig_5) (*Figure 5A* and *5B*).^69–79^

**Figure 5.**
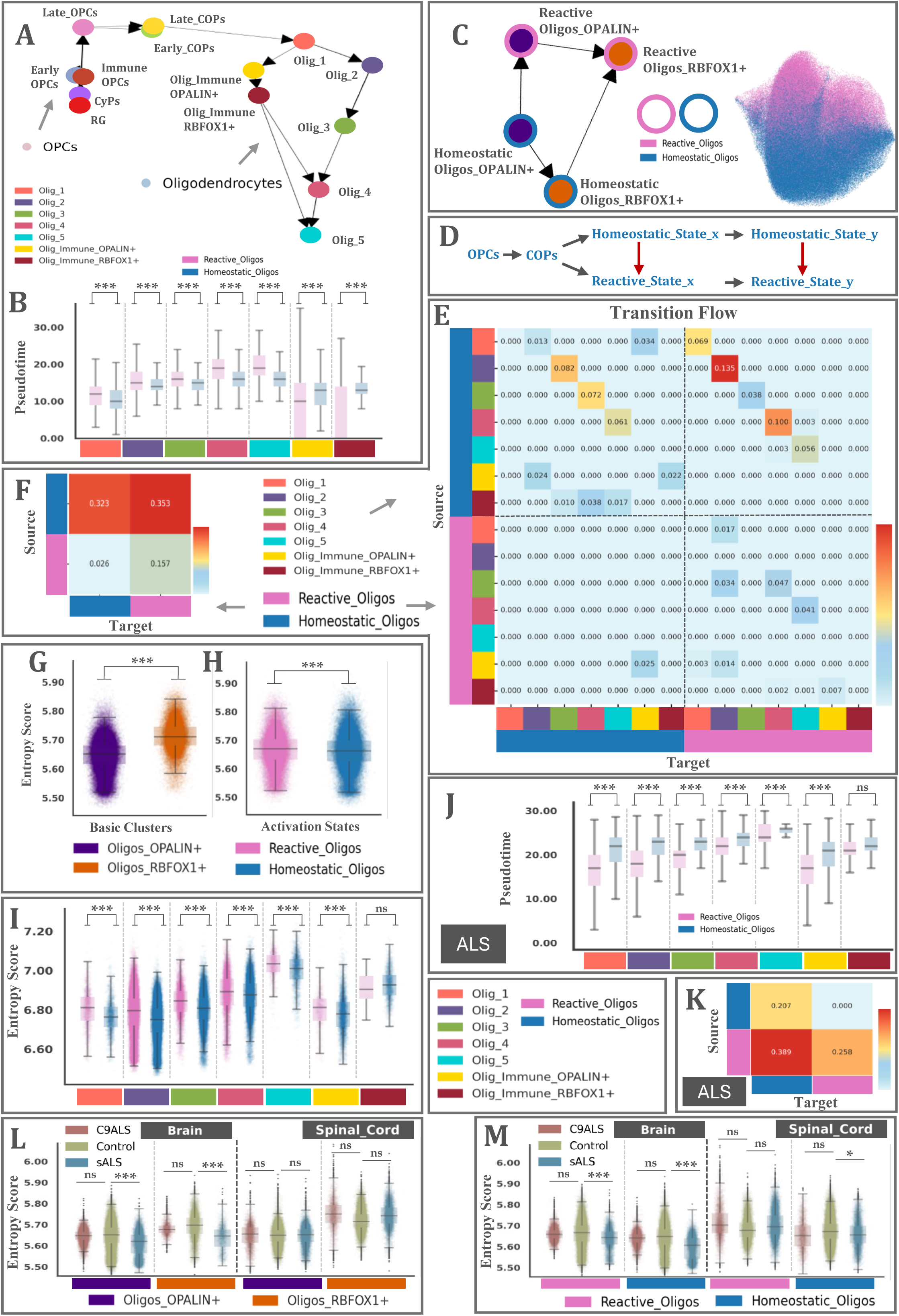
Trajectories and Transition Probabilities of Oligodendrocyte Activation States. **(A)** Abstract plot of the maturation trajectory between subclusters of the motor cortex and spinal cord control OPCs and oligodendrocytes. Each circle represents a subcluster. **(B)** Boxplots of the pseudotemporal ordering by subcluster grided by activation state in the control motor cortex and spinal cord. For each subcluster there is one box for the reactive state (pink) and one for the homeostatic (blue). Significance is calculated by a linear mixed-effects model. **(C)** Abstract plot of the maturation and activation trajectory between basic clusters grouped by activation states of the motor cortex and spinal cord control oligodendrocytes. Each circle in the oligodendrocyte population represents a basic cluster grouped by activation state. **(D)** Representation of the suggested maturation and activation trajectory between activation states. **(E)** Matrix plot of the transition probabilities between each oligodendrocyte subcluster grouped by activation state in the control motor cortex and spinal cord. **(F)** Matrix plot of the transition probabilities aggregated at a coarse level between each activation state in the control motor cortex and spinal cord. **(G)** Boxplots of the calculated cell-level entropy grouped by basic cluster, Oligos_OPALIN+ (purple) and Oligos_RBFOX1+ (orange) in the control motor cortex and spinal cord. Significance is calculated by a linear mixed-effects model. **(H)** Boxplots of the calculated cell-level entropy grouped by activation state, reactive (pink) and homeostatic (blue) in the control motor cortex and spinal cord. Significance is calculated by a linear mixed-effects model. **(I)** Boxplots of the calculated cell-level entropy by subcluster grided by activation state. For each subcluster there is one box for the reactive state (pink) and one for the homeostatic (blue). Significance is calculated by a linear mixed-effects model. **(J)** Boxplots of the pseudotemporal ordering by subcluster grided by activation state in the ALS (C9ALS and sALS) spinal cord. For each subcluster there is one box for the reactive state (pink) and one for the homeostatic (blue). Significance is calculated by a linear mixed-effects model. Significance is calculated by a linear mixed-effects model. **(K)** Matrix plot of the transition probabilities aggregated at a coarse level between each activation state in the ALS (C9ALS and sALS) spinal cord. **(L – M)** Boxplots of the calculated cell-level entropy by basic cluster **(L)** and activation state **(M)** grided by status. For each by basic cluster (L) and activation state (M) there is one box for each status (C9ALS, Control, sALS). Significance is calculated by a linear mixed-effects model. Abbreviations in this figure include: lmm; Linear mixed-effects model lmm * p-value < 0.05, ** p-value < 0.01, *** p-value < 0.001 See also **Figure S3**.

Multiple trajectory inference methods were independently applied, including Slingshot^71^, Monocle3^72–75^, Lamian^76^, Palantir^77^, PAGA^78^, and MARGARET^79^ to ensure the robustness of inferred developmental paths. For clarity, the figures primarily display results from MARGARET, which provides probabilistic modeling of latent time and gene program transitions. All trajectory patterns were cross-validated across the different methods, yielding consistent results.

Next, we leveraged MARGARET and PAGA to generate a global topological view of oligodendrocyte subclusters, focusing on the relationships between reactive and homeostatic activation states.^78,79^ In this abstract graph, every node represents a stable or metastable state (e.g., Olig_1, Olig_5) and each directed edge represents a predicted transition. Branch transition probabilities calculated through the MARGARET pipeline indicated that transitions predominantly occur from homeostatic to reactive states, rather than the reverse (*Figure 5C-5F*). Homeostatic cells can progress along the homeostatic trajectory (e.g., Homeostatic_Olig_1 → Homeostatic_Olig_2) or laterally transition to the corresponding reactive state (e.g., Homeostatic_Olig_1 → Reactive_Olig_1). Reactive cells, in contrast, mostly progressed forward along the reactive trajectory, with minimal evidence for reversion to homeostatic states.

To further characterize the directionality of oligodendrocyte state transitions, we quantified the pseudotime-oriented transition flow between subclusters grouped by activation state (*Figure 5E-5F*). Using the mean pseudotime per cluster, we retained only edges connecting clusters with increasing pseudotime (source < target), thus capturing transitions that follow the inferred biological trajectory. We then aggregated these directed edges at the state level. The resulting global transition flow matrix was normalized by the total transition weight. This approach revealed a clear asymmetry: transitions from homeostatic to reactive oligodendrocyte states dominated over reverse transitions, consistent with a progressive activation process along pseudotime (*Figure 5E*). When aggregated at a coarse level, the overall transition probability from homeostatic to reactive states exceeded the inverse direction, supporting the theoretical expectation of a unidirectional shift toward reactivity (*Figure 5F*).

To evaluate the robustness of this directional bias, we performed both per-donor and bootstrap resampling analyses. For the per-donor analysis, transition flows were computed independently for each donor, allowing us to account for inter-individual variability. Across donors, the homeostatic → reactive flow remained consistently higher than the reactive → homeostatic flow, as confirmed by a linear mixed-effects model [lmm] (p-value < 0.001). In addition, bootstrap resampling of cells (1,000 iterations) yielded a mean directional difference of 0.397, with a 95% confidence interval for the net transition difference (H→R − R→H). The confidence interval did not overlap zero, indicating a statistically significant and highly stable directional imbalance. Together, these analyses demonstrate that the observed pseudotime-oriented transition from homeostatic to reactive states is both biologically consistent and statistically robust.

We subsequently quantified cell-level entropy across various datasets and tissue types (*Figure 5G-5I*).^77,83–85^ Because entropy reflects system disorganization and dynamic change, it serves as a proxy for cellular instability. Linear mixed-effects modeling revealed significant differences in scEntropy among basic clusters, with Oligos_RBFOX1+ displaying higher entropy compared to Oligos_OPALIN+ both in the brain (β = 0.092, SE < 0.001, p-value < 0.001) and in the spinal cord (β = 0.131, SE < 0.001, p-value < 0.001) (*Figure 5G*). The random intercept for sample showed negligible variance (σ² ≈ 0), indicating minimal sample-to-sample variability in entropy. Likewise, reactive cell states consistently exhibited elevated scEntropy relative to homeostatic states (*Figure 5H*). Although the effect size was smaller (brain: β = 0.047, SE = 0.001, p-value < 0.001 and spinal cord: β = 0.037, SE = 0.001, p-value < 0.001), the association remained highly robust, and sample-level variance was again negligible (σ² ≈ 0), suggesting that entropy variation primarily arises within rather than between samples. These differences in the activation state entropy were conserved in the subcluster level (*Figure 5I*).

To study transcriptional dynamics, we applied RNA velocity analysis. This approach quantifies the unspliced-to-spliced transcript ratios across subclusters.^80–81^ Homeostatic populations exhibited higher unspliced transcript ratios (lmm; p-value < 0.001), suggesting a less mature, more plastic state poised for transition (*Figure S3A*). We also inferred splicing kinetics and reconstructed a continuous vector field of cell-state transitions, enabling prediction of future cell fates, using Dynamo^82^ (*Figure S3B*). This extends RNA velocity to estimate the most probable future states of cells based on transcriptional dynamics. A vector field represents each cell’s state as a point in a low-dimensional gene-expression landscape, with an arrow showing the direction and speed of its likely transcriptional change. In Dynamo, these arrows are derived from RNA velocity and reconstructed into a continuous, smooth field that predicts how any cell in the landscape will move and what states it may reach. In this model, “movement” reflects changes in gene expression over time rather than physical motion.

However, interpretations of such analyses must be made with caution, as they rely on temporal inference from inherently static, snapshot transcriptomic data. Within this limitation, we propose a quantitative interpretation of the single-cell potential landscape between activation states as inferred from our data.

To do so, we computed the single-cell potential using Dynamo’s vf.potential() function. In this context, “potential” does not represent physical energy but a numerical approximation of system stability derived from the reconstructed vector field.^86–88^ Dynamo estimates potential based on the negative log of local cell density within the learned dynamical manifold, providing a quantitative measure of how likely or stable each transcriptional state is. Cells located in high-density regions correspond to low-potential (stable attractor basins), whereas those in sparse regions correspond to high potential (transient or unstable states). Importantly, Dynamo reorients potential values (high = stable, low = transient) to mimic the inferred pseudotime direction, so cells appear to ‘flow’ from low- to high-potential regions along activation or maturation trajectories.

Oligos_RBFOX1+ and homeostatic cells appeared to have higher single-cell potential (residing in basins) than Oligos_OPALIN+ and reactive cells (located at the tops of hills) (*Figure S3C*). Focusing on the differences between activation states, mean single-cell potential (μ) differed significantly between activation states grouped by basic clusters (lmm, p-value < 0.001). Homeostatic oligodendrocytes occupied higher-potential basins. Homeostatic_Oligos_OPALIN+ (μ = 1.01 ± 0.87) and Homeostatic_Oligos_RBFOX1+ (μ = 4.97 ± 1.23). Reactive cells resided in lower-potential, transient regions: Reactive_Oligos_OPALIN+ (μ = 0.68 ± 0.61) and Reactive_Oligos_RBFOX1+ (μ = 3.95 ± 1.09). Relative to their reactive counterparts, homeostatic clusters displayed significantly elevated potential (reactive vs. homeostatic Oligos_OPALIN+: β = +0.32, SE = 0.02, p-value < 0.001; reactive vs. homeostatic Oligos_RBFOX1+: β = +1.02, SE = 0.03, p-value < 0.001). This quantitative asymmetry in the reconstructed energy landscape supports the interpretation of homeostatic states as stable attractors and reactive states as dynamically unstable transition zones.

Supporting vector-field metrics, such as divergence, computed directly from the reconstructed field, further reinforced this interpretation. Divergence measures whether the arrows (vectors) around a point converge or diverge. Positive divergence indicates that arrows point outward, suggesting an unstable or transient state. Negative divergence indicates convergence, corresponding to an attractor state, such as a differentiated cell type. Reactive and Oligos_OPALIN+ populations displayed higher divergence, whereas homeostatic states and Oligos_RBFOX1+ populations exhibited lower (lmm; p-value < 0.001) (*Figure S3D*). These orthogonal descriptors converge on the same interpretation as single-cell potential, reinforcing the idea that reactive states occupy higher single-cell potential, unstable regions. This framework outlines a continuum in oligodendrocytes. Stable, yet plastic, multipotent homeostatic states occupy low-potential valleys of the cellular landscape. Reactive states reside in high-potential, unstable regions, where external stimuli can push cells toward a bifurcation point, deciding whether to remain homeostatic or transition into reactivity.

### Altered Oligodendroglial Trajectories and Entropy in ALS

Topographic lineage mapping further showed that in both ALS and control samples, the initial post-OPC oligodendrocyte population is either homeostatic or reactive Olig_1, suggesting a conserved early lineage trajectory across conditions. Dynamic trajectory analysis and lineage inference in ALS spinal cord samples revealed a reversal in pseudotime progression compared to controls (*Figure 5B*): in ALS, homeostatic states exhibited significantly higher pseudotime values than reactive states (*Figure 5J*). Consequently, in the disease condition, the pseudotime-oriented transition flow showed a predominant direction from reactive toward homeostatic oligodendrocyte states (lmm; p-value < 0.001), indicating a reversal of the activation trajectory, where cells tend to revert towards homeostasis (*Figure 5K*). Both per-donor linear mixed-effects modeling and bootstrap resampling (1,000 iterations) confirmed a statistically significant reversed directional bias, as the 95% confidence interval for the mean transition difference did not include zero. Transcriptional entropy was significantly reduced in the sALS motor cortex compared to the controls, and in homeostatic sALS spinal cord samples (*Figure 5L-5M*).

Consistent with findings in controls, single-cell potential remained higher in homeostatic states than in reactive states within ALS samples, placing them in low-potential basins of the single-cell potential landscape (*Figure S3E-S3F*).

RNA velocity analyses were performed exclusively on our in-house datasets, as these were the only datasets containing sufficient spliced and unspliced read information. In the motor cortex, some populations—particularly OLIGOS_RBFOX1+ cells—had insufficient read coverage to support reliable RNA velocity estimation.

### Gene Regulatory Network Analysis Reveals Distinct Transcriptional Control of Activation States in ALS

To explore the regulatory mechanisms underlying transitions between activation states, we constructed gene regulatory networks (GRNs) using CellOracle and SCENIC, across the motor cortex and spinal cord.^89–90^ SCENIC was employed to infer transcription factor (TF)–target relationships based on co-expression and cis-regulatory motif enrichment, defining regulons that represent putative, direct regulatory interactions. In parallel, CellOracle integrated prior TF–target annotations with our single-cell transcriptomic data to reconstruct condition-specific GRNs, thereby capturing context-dependent regulatory connectivity within each activation state. In an initial integrative analysis, we focused on control conditions, identifying a core set of transcription factors (TFs) conserved across both motor cortex and spinal cord that correlated with the shift between reactive and homeostatic cellular states. This analysis aimed to find specific TF sets that either promote the activation of reactive gene programs or actively suppress them to maintain cellular and tissue homeostasis. We reconstructed GRNs across both reactive and homeostatic cell states and quantified collective transcription factor activity per activation state.

From the CellOracle-inferred regulatory network, the transcription factors (TFs) exhibiting the highest degree centrality (reflecting the number of direct regulatory connections) included *TCF12*, *MEF2A*, *PBX3*, *NR3C1*, *STAT2*, *NPAS3*, *KLF7*, *ELF1*, *FOXN2*, *ZBTB7A*, *CHD2*, *RAD21*, *TCF7L2*, *JUND*, *KLF12*, *MAX*, *NRF1* and *ZNF148*. In terms of betweenness centrality, which quantifies how often a node lies on the shortest paths between other nodes, reflecting its role as a “bridge” in the network, the top-ranking TFs were *NPAS3*, *TCF12*, *HIVEP3*, *PBX3*, *PBX1*, *STAT2*, *KLF12*, *CREB5*, *JUND*, *ZBTB16*, *MAX*, *ATF7*, *GTF2I*, *KLF9*, *NKX6-2*, *BBX*, *NF1*. These TFs likely represent central regulators within the overall transcriptional network. To identify which of them may specifically govern transcriptional programs associated with cellular activation, we further refined the network by retaining only edges connected to genes differentially expressed between activation states under control conditions. In this filtered network, the TFs with the highest degree and betweenness centrality were *TCF12*, *RAD21*, *JUND*, *NR3C1*, *BCLAF1*, *PBX3*, *NRF1*, *ELF1*, *FOXK2*, *VEZF1* and *NPAS3*, *JUND*, *NKX6-2,* respectively (*Figure 6A*).

**Figure 6.**
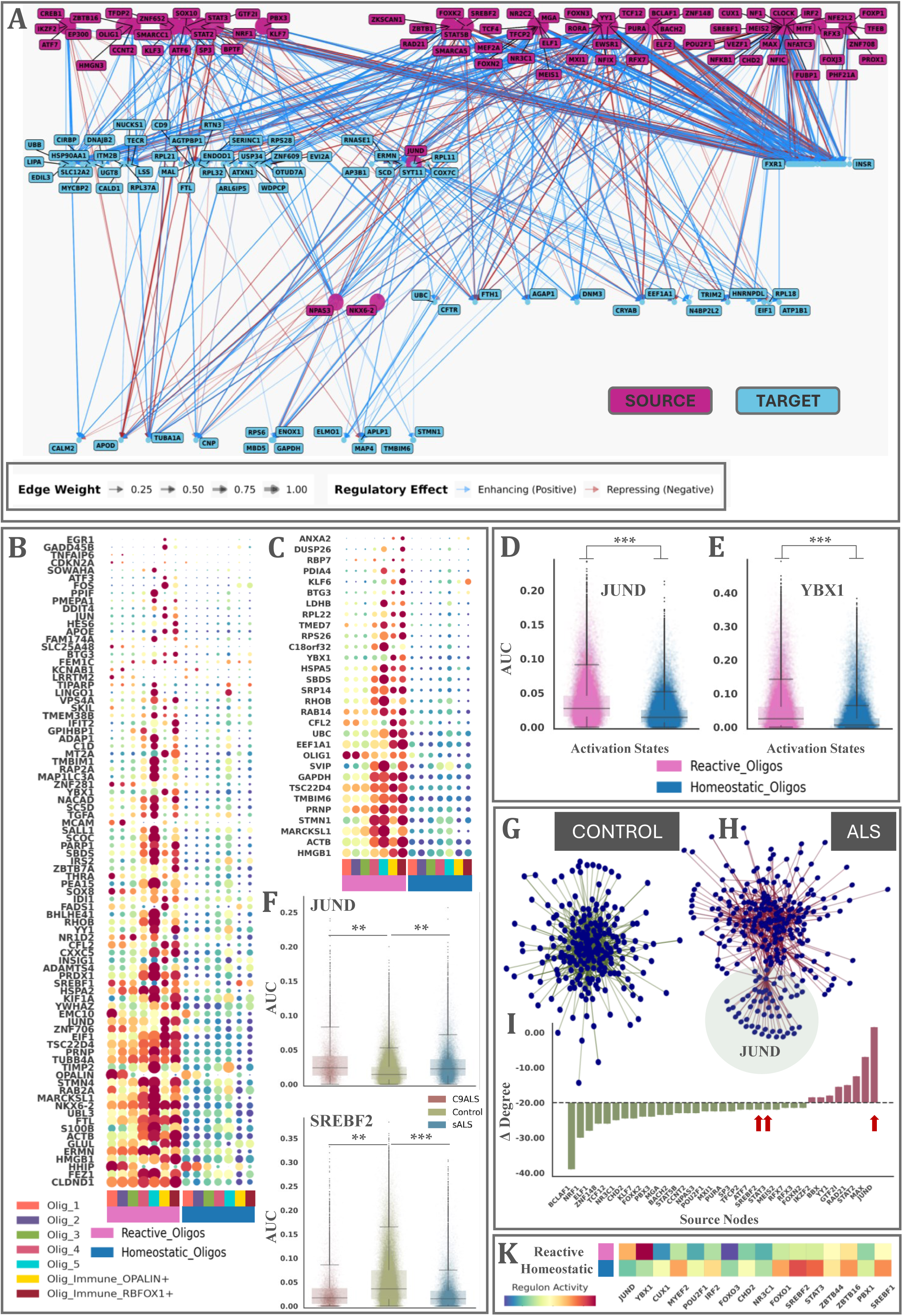
JUND as a Key Regulatory Factor for Transition Between Activation States. **(A)** GRN graph of the main TFs and their target genes in the control motor cortex and spinal cord oligodendrocytes. The nodes represented are the top ones selected based on centrality metrics. **(B)** DotPlot of the JUND regulon (SCENIC). Each column is a subcluster grouped by activation state. **(C)** DotPlot of the YBX1 regulon (SCENIC). Each column is a subcluster grouped by activation state. **(D - E)** Boxplots of the calculated AUC score (SCENIC) of the JUND **(D)** and the YBX1 **(E)** regulons grouped by activation state, reactive (pink) and homeostatic (blue) in the control motor cortex and spinal cord. Significance is calculated by a linear mixed-effects model. **(F)** Boxplots of the calculated AUC score (SCENIC) of the JUND **(D)** and the SREBF2 **(E)** regulons grouped by status, C9ALS (pink), control (green) and sALS (blue) in the spinal cord. Significance is calculated by a linear mixed-effects model. **(G - H)** GRN graphs of the main TFs and their target genes in the control motor cortex and spinal cord **(G)** and the ALS spinal cord **(H)** oligodendrocytes. The nodes represented are the target genes differentially expressed between activation states and their respective source nodes (TFs). **(I)** Sorted bar chart of degree changes ((Δ Degree) for source nodes in ALS compared to the control. Red increased degree centrality in the disease and green decrease degree centrality in the disease. **(K)** Matrix plot showing the AUC values of SCENIC-derived regulons, grouped by activation state. Abbreviations in this figure include: AUC; Area Under the Curve, lmm; Linear mixed-effects model lmm * p-value < 0.05, ** p-value < 0.01, *** p-value < 0.001

We further investigated GRNs using SCENIC, identifying 21 regulons enriched within the oligodendrocyte population. Among these, only the JUND and YBX1 regulons were linked to target genes that were differentially expressed between activation states (*Figure 6B-6C* and *6K*). Both regulons exhibited significant enrichment in reactive oligodendrocyte states, as determined by linear mixed models (*JUND*: β = 0.0246, SE < 0.001, p < 0.001; *YBX1*: β = 0.0452, SE < 0.001, p < 0.001) (*Figure 6D-6E*). Nevertheless, YBX1 ranked low in both degree and betweenness centrality within the CellOracle-derived network, suggesting a more peripheral regulatory role despite its differential activity.

We further extended our GRN reconstruction to investigate disease-associated network alterations in both the motor cortex and spinal cord. Our analysis specifically focused on the spinal cord, where previous results indicated a shift toward reactive states, with the aim of identifying regulons that may drive this transition. Among all regulons identified by SCENIC analysis, the *JUND* regulon exhibited increased activity in the ALS spinal cord (both C9ALS and sALS: p-value < 0.01), as well as the *ZBTB44* regulon (C9ALS p-value < 0.05; sALS p-value < 0.001) (*Figure 6F*). Consistent with our network analysis, *JUND* target genes were differentially expressed in reactive compared to homeostatic oligodendrocyte states, while *ZBTB44* target genes showed differential expression between early and late maturation oligodendrocytes. These patterns further support the enrichment of reactive and early post-OPC populations in the ALS spinal cord. In contrast, *SREBF2* (C9ALS p-value < 0.01; sALS p-value < 0.001) showed decreased activity in disease and the *STAT3* regulon was significantly downregulated specifically in the sALS spinal cord (p-value < 0.01) (*Figure 6F*). All other regulons did not exhibit significant changes in activity under disease conditions. Within the CellOracle-constructed networks, *JUND* emerged as the top node with increased degree centrality — a measure of how many direct connections a node has in the network and thus its potential influence — while *SREBF2* and *STAT3* showed decreases in degree centrality, but they were not among the top hits (*Figure 6I*).

In the motor cortex, changes in regulon activity were observed only in the sALS condition, with activation of regulons associated with homeostatic states (*Figure 6K*), including *MYEF2*, *POU2F1*, *IRF2*, *FOXO3*, *CHD2*, *NR3C1*, *FOXO1*, *ZBTB44*, and *SREBF1*. Notably, *JUND* activity was unchanged in the diseased motor cortex, whereas *ZBTB44* was enriched.

Interestingly, recent studies have implicated *JUND* in driving the transition of astrocytes into a reactive state, and it has also been identified as a master regulator of gene expression in oligodendrocytes.^91,92^

### Metabolic Activity and Inter/Intra-Lineage Communication in Control and ALS

Beyond their well-established role in myelination, oligodendrocytes also support neurons by providing metabolic resources and contributing to immune regulation.^11,13^ They achieve this through active communication with both neuronal and glial cells. To better understand these additional roles—particularly how metabolism and intercellular communication vary across different oligodendrocyte states—we systematically analyzed the metabolic profiles and cell–cell communication (CCC) networks of the two identified activation states.^93–97^ We then assessed their interactions not only with other brain cell types but also within the oligodendrocyte lineage itself.

We leveraged scMetabolism to infer pathway-level metabolic differences from transcriptional pathway expression between the activation states.^93^ The two groups showed distinct metabolic profiles (*Figure 7A*). Reactive oligodendrocytes displayed broad metabolic activation, with enrichment in glucose, glycogen, fatty acid, glutamate, and cofactor metabolism^98^, alongside PKA-signaling^99^, nitric oxide production^100^, and arachidonic acid metabolism^101^—pathways linked to inflammatory lipid mediators and stress responses (*Figure 7A*). In agreement with our previous analysis, this pattern suggests a reactive oligodendrocyte state involved in rapid energy mobilization and lipid remodeling. In contrast, homeostatic oligodendrocytes were enriched in lipid-centered homeostatic pathways, including PPARα-regulated fatty acid oxidation^102^, carnitine metabolism^103^, biotin-dependent fatty acid synthesis^104^, and glycosaminoglycan metabolism^105–107^, consistent with homeostatic cells maintaining long-term lipid balance and extracellular matrix integrity (*Figure 7A*). Previous studies have also shown that glycosaminoglycans may act as inhibitors of OPC and oligodendrocyte differentiation.^105–107^

**Figure 7.**
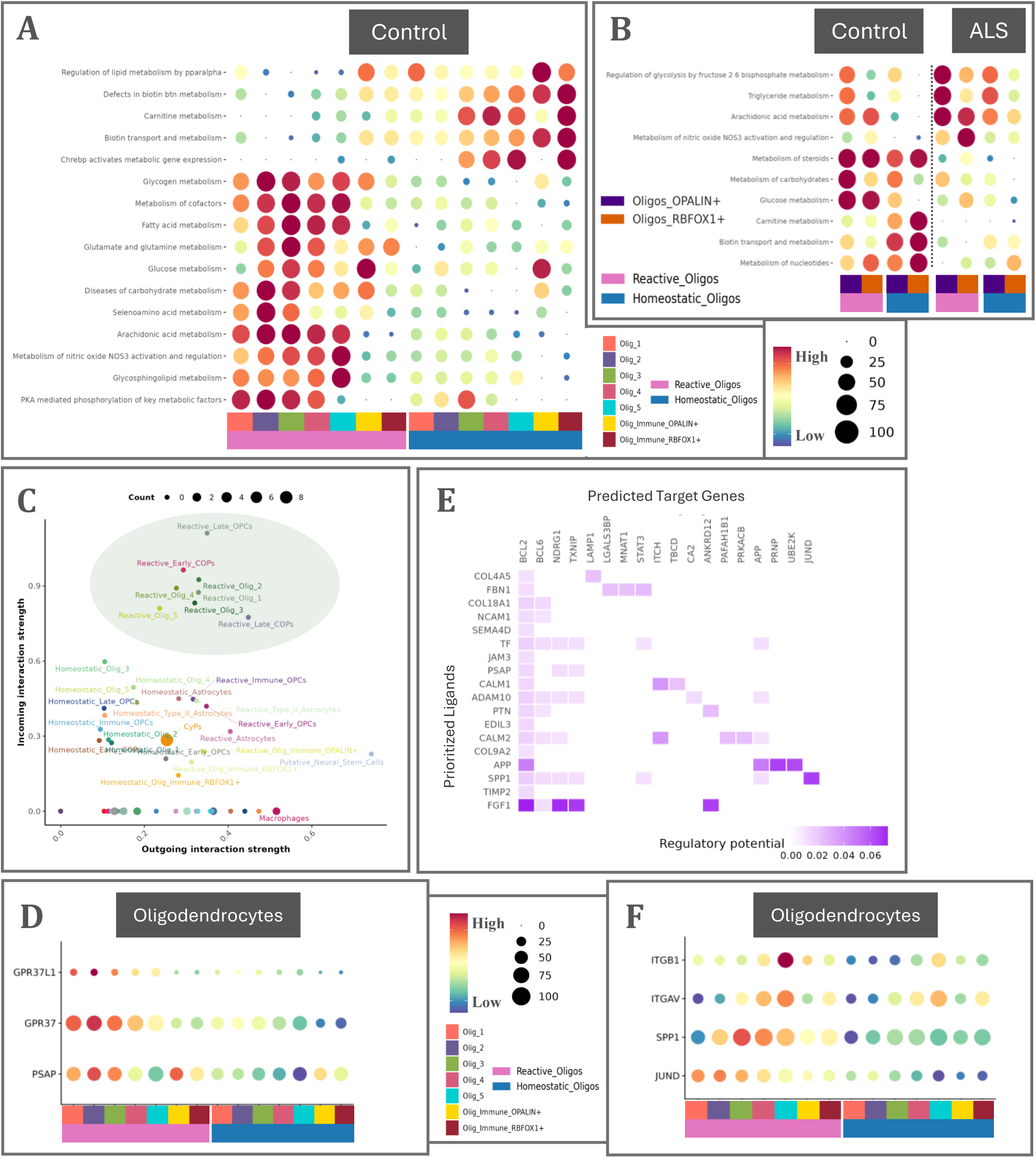
Metabolic and Communication Profile in Control and ALS. **(A)** DotPlot of metabolic pathways (REACTOME) being differentialy expressed between reactive and homeostatic subcluster states in motor cortex and spinal cord control oligodendrocytes. **(B)** DotPlot of metabolic pathways (REACTOME) being differentialy expressed between control and ALS (C9ALS and sALS) spinal cord oligodendrocytes, grouped by basic clusters and activation states. **(C)** ScaterPlot of the PSAP signaling network in the motor cortex and spinal cord control subclusters and activation states. **(D)** DotPlot of the PSAP signaling network expression in motor cortex and spinal cord control oligodendrocytes, grouped by subclusters and activation States. **(E)** HeatMap of the prioritized intra-oligodendrocyte population ligands and their predicted target genes. **(F)** DotPlot of the SPP1 signaling network expression in motor cortex and spinal cord control oligodendrocytes, grouped by subclusters and activation States

Next, we investigated the metabolic differences between controls and ALS. In the spinal cord, the control population exhibited enrichment in nucleotide, carbohydrate, glucose, and steroid metabolism, along with biotin and carnitine pathways—suggesting a profile oriented toward anabolic activity, energy production, and biosynthesis, supporting cell growth and maintenance (*Figure 7B*). In contrast, the ALS population (C9ALS and sALS) demonstrated enrichment in nitric oxide production, arachidonic acid metabolism, triglyceride metabolism, and glycolytic regulation by fructose 2,6-bisphosphate—indicating a stress-responsive, inflammatory, and energy-mobilizing state, with pathways tied to signaling lipids, vascular modulation, and rapid energy release (*Figure 7B*).

In the motor cortex, metabolic alterations were less pronounced but more diverse. The sALS population was enriched in glucose, glycogen, and carbohydrate metabolism, thereby resembling the control population and suggesting a carbohydrate-centered energy strategy, showing rapid ATP production through glycolysis and glycogen breakdown. This profile suggests cells primed for immediate energy demands and metabolic flexibility. In the C9ALS population, we observed downregulation of pathways that were enriched in the sALS population, particularly those related to triglyceride metabolism and PKA-mediated phosphorylation. This molecular profile closely resembles that seen in spinal cord ALS. This reflects a lipid-based energy strategy coupled with hormonal and signaling regulation of metabolism, indicating reliance on stored fats and dynamic metabolic control—often seen in longer-term energy mobilization or signaling-driven metabolic adaptation.

We leveraged CellChat and LIANA+ to profile CCC networks across cell types.^94–96^ Among the analyzed signaling pathways, *PSAP* (prosaposin) signaling emerged as one of the most differentially regulated between activation states (*Figure 7C*). Reactive states exhibited enhanced *PSAP*-mediated communication (*Figure 7C*). Both activation states functioned as major senders and receivers of PSAP signals, engaging in interactions with multiple cell types, including astrocytes (both reactive and homeostatic), microglia (disease-associated and homeostatic), macrophages, vascular cells, and neurons. However, the reactive state exhibited markedly higher capacity to receive PSAP signaling (*Figure 7C*).

The receptors for PSAP, GPR37 and GPR37L1, were differentially expressed across oligodendrocyte lineage cells: GPR37 was enriched in oligodendrocytes, while GPR37L1 was enriched in OPCs (*Figure 7D*). Both receptor transcripts were upregulated in the reactive state compared to the homeostatic state, consistent with the increased signal reception observed in the communication analysis (*Figure 7D*). This expression pattern remained largely unchanged under disease conditions.

Previous studies indicated that GPR37 signaling exerts dual effects on myelination: its activation can negatively regulate myelin formation, whereas loss of GPR37 reduces myelin-associated glycoprotein (MAG) production.^108–109^ This suggests that GPR37 may act as a bidirectional regulator of myelin synthesis. In our dataset, its increased expression in reactive oligodendrocytes—cells actively engaged in myelination—may represent a negative feedback mechanism to fine-tune myelin production and prevent over-myelination.

We next analyzed intra-lineage communication among oligodendrocytes. Using ligand-receptor interaction analysis, the top prioritized ligands—including PSAP, TF, CALM2, CALM1, SPP1—were selected for downstream analysis using NicheNet to predict their potential downstream target genes.^97^ A key finding was the consistent prediction of *BCL2* as a common target across these ligands (*Figure 7E*). *BCL2*, a well-characterized anti-apoptotic gene, was upregulated in homeostatic and immune-related oligodendrocytes, suggesting a potential role in promoting cell survival and stability.^110^ Importantly, our analysis identified *JUND* as the top target gene of the SPP1 ligand (*Figure 7E-7F*). *SPP1*, previously implicated in cellular reactivity, was enriched in reactive oligodendrocytes in our data.^111–112^

Finally, cell–cell signaling did not show significant alterations in any ALS group, suggesting that these communication processes may remain largely preserved or tightly regulated during ALS progression.

## DISCUSSION

Our study reveals a previously unappreciated spectrum of oligodendrocyte activation states in the human motor system, highlighting how homeostatic and reactive states dynamically balance in health and disease. By analyzing more than one million single nuclei—including *350,000* oligodendrocyte and oligodendrocyte precursor (OPC) cells—from 176 postmortem samples, we establish a simple conceptual framework that captures the essential features of this system’s behavior.

We propose that oligodendrocytes predominantly exist within two principal activation states—homeostatic and reactive—that define poles of a continuous transcriptional spectrum rather than discrete subtypes. Even in control tissue, homeostatic oligodendrocytes exhibit low-level reactivity, suggesting that this axis reflects an intrinsic adaptive capacity rather than a purely pathological feature. Regionally, the spinal cord contains a larger proportion of homeostatic cells, whereas the motor cortex exhibits a more balanced distribution between the two states, potentially reflecting the higher plasticity of cortical circuits. In ALS, this equilibrium shifts toward reactivity, indicating that system-level stress promotes activation of stress-associated programs. Interestingly, the inferred transition probabilities suggest enhanced re-entry from reactive to homeostatic states in ALS, consistent with compensatory attempts to restore equilibrium under stress.

High-resolution transcriptomic profiling identifies novel subclusters and markers, including Olig_5 and *PPEF1*, and highlights *LRP2* as an oligodendrocyte-enriched gene that may serve as a molecular indicator of lineage identity. These findings provide molecular entry points for dissecting oligodendrocyte heterogeneity and lineage plasticity. The compositional and transcriptional differences observed between the motor cortex and spinal cord—and their shifts in ALS—underscore the region- and disease-specific nature of oligodendrocyte activation. Elevated CSF levels of MBP and LRP2, together with the enrichment of stress- and death-related pathways in reactive states, are consistent with myelin disruption and cell vulnerability, although we cannot exclude alternative mechanisms such as extracellular vesicle release contributing to these signals.

Our integrative analysis of transcriptional regulation identifies *JUND* as a transcription factor associated with reactive oligodendrocyte states. In contrast, downregulation of *SREBF2* and *STAT3* may modulate these transitions. Although we do not claim direct causality, these associations suggest testable hypotheses regarding the transcriptional control of glial activation. The co-occurrence of *JUND* upregulation with increased *FTH1*/*FTL* expression aligns with a coordinated stress-response module linking iron homeostasis and cellular resilience, consistent with known *JUND*-dependent regulation of *FTH1* in non-CNS systems.^113^ Future perturbation studies—using human-derived model systems or in vivo manipulations—will be necessary to establish whether these transcriptional associations reflect causal regulatory relationships.

Beyond transcriptional control, we observed distinct metabolic and intercellular communication programs across oligodendrocyte states. Reactive cells upregulated glycolytic and lipid-stress pathways, consistent with increased energetic demand and stress vulnerability, whereas homeostatic states favored lipid maintenance and survival-associated processes. In ALS, this balance shifted further toward metabolic stress signatures, in line with impaired trophic support. Notably, reactive states engaged more extensively in multicellular signaling, including prosaposin–GPR37/37L1 interactions with astrocytes, microglia, and neurons. These ligand–receptor changes converged on transcriptional regulators such as *JUND*, suggesting feedback between intracellular stress responses and intercellular communication. Together, these observations imply that oligodendrocyte activation is not purely cell-autonomous but emerges from coordinated crosstalk within neural networks.

The homeostatic-to-reactive continuum described here may represent a general organizing principle of CNS glia, paralleling activation spectra reported for astrocytes and microglia. The conservation of these transcriptional programs across neurodegenerative and psychiatric conditions, as well as solid tumors,^114^ suggests that glial reactivity follows shared regulatory logic tuned by context and region. Expanding this framework through multi-condition, multi-region single-cell atlases will be crucial for distinguishing universal from disease-specific mechanisms and for determining whether reactive states are protective adaptations or maladaptive endpoints.

By applying a single-cell potential landscape framework, we propose that homeostatic oligodendrocytes occupy stable attractor states, whereas reactive cells populate transient, higher-potential regions poised for adaptive responses. We emphasize that vector field and trajectory inferences should be interpreted with caution, as transcriptomic data alone cannot resolve temporal causality. Nonetheless, these analyses offer a useful abstraction for understanding population-level stability and transitions. This continuum challenges earlier categorical models of “disease-associated oligodendrocytes” (DAOs) and instead supports a view of glial activation as a fluid, context-dependent process.

While the continuous nature of activation complicates subtype classification, it reinforces the utility of dynamic frameworks to capture cellular plasticity. Transcriptomic inference of metabolic and signaling programs, though indirect, provides a roadmap for targeted functional validation through integrated spatial transcriptomics, proteomics, and imaging approaches. Exploring how intrinsic transcriptional regulators interact with extrinsic cues to govern state transitions remains an important future goal.

In summary, our study defines a comprehensive atlas of oligodendrocyte heterogeneity and activation dynamics in the human motor system, uncovering novel markers, candidate regulators, and principles of glial state equilibrium. We propose that homeostatic oligodendrocytes reside in low-potential basins functioning as reservoirs for adaptive activation, whereas sustained reactivity—particularly in ALS—drives an accumulation of metabolically demanding, vulnerable states that may contribute to cell loss and myelin dysfunction. By framing oligodendrocyte behavior as a continuum of activation rather than a dichotomy of health and disease, our work establishes a conceptual foundation for future studies aimed at modulating glial states to preserve neural circuit integrity and counteract neurodegeneration.

## Methods

### RESOURCE AVAILABILITY

#### Lead Contact

Further information and requests for resources, data, or code should be directed to and will be fulfilled by the Lead Contact, **Maria Georgopoulou and Philip Van Damme.**

#### Materials Availability

This study did not generate new unique reagents.

#### Data and Code Availability

- Processed AnnData (.h5ad) and Seurat (.rds) objects generated during this study are available online in Zenodo at https://doi.org/10.5281/zenodo.17669249.

Public datasets used in this work are listed in **Methods (*Public Data Acquisition)*** with corresponding accession numbers.ffi

- Custom scripts and notebooks for preprocessing, integration, and visualization are available at https://github.com/mariageorgopoulou/Oligodendrocyte_Transcriptomic_Atlas_2025/tree/main.
- Temporary restrictions apply to data availability.
- No restrictions apply to code availability.

### EXPERIMENTAL MODEL AND SUBJECT DETAILS

#### In-house Sample Collection, Preparation and Single-nucleus RNA sequencing

Postmortem human motor cortex and spinal cord tissue were obtained from clinically and neuropathologically confirmed cases of ALS and age-matched controls. The cohort included 15 samples per tissue across three conditions—five control, five *C9orf72*-ALS (C9ALS), and five sporadic ALS (sALS)—yielding a total of 30 samples. Information on the in-house postmortem human samples collection, preparation and sequencing can be found in the Masrori, P., Bijnens, B., Fumagalli, L. et. al, Nat Neurosci 28, 2217–2230 (2025). https://doi.org/10.1038/s41593-025-02075-1.

### METHOD DETAILS

#### Single-nucleus RNA Sequencing (snRNA-seq)

Single nuclei were isolated from frozen postmortem tissue following previously described protocols (Masrori, P., Bijnens, B., Fumagalli, L. et. al, Nat Neurosci 28, 2217–2230 (2025). https://doi.org/10.1038/s41593-025-02075-1). Libraries were generated using the Chromium Single Cell 3′ v3 kit (10x Genomics) and sequenced on the NovaSeq 6000 platform.

#### Demultiplexing of Pooled Sequencing Runs

Sequencing runs were demultiplexed using Freemuxlet. Sample barcodes were extracted from the _freemuxlet.clust1.samples.gz output and merged with Cell Ranger-generated raw count matrices. Seurat objects were created using the CreateSeuratObject() function and annotated with Freemuxlet metadata (AddMetaData()). Only singlets (DROPLET.TYPE == “SNG”) with a confident genotype assignment (BEST.GUESS == “1,1”) were retained. Each sample was assigned a Sample_ID and saved as an .rds file for downstream analysis.

Filtered singlet Seurat objects were converted to AnnData format using the as.anndata() function from *SeuratWrappers*, retaining the raw count matrix as an additional layer. Python integration was performed through *reticulate*, specifying the Python 3.9 environment containing the anndata module. Converted files were stored as .h5ad objects for preprocessing in Python.

#### Public Data Acquisition

We collected public snRNA-seq data from 15 studies (16 datasets). Original data can be found in GEO (https://www.ncbi.nlm.nih.gov/geo/) with accession numbers: **GSE219280**, **GSE226753**, **GSE271156**, **GSE229169**, **GSE228778**, **GSE190442**, **GSE287257, GSE249210, GSE171892**, and **GSE243077.** The rest from the data can be acquired from: https://www.synapse.org/Synapse:syn51105515/files/(syn51105515), https://assets.nemoarchive.org/dat-ek5dbmu(RRID:SCR_016152), https://www.ncbi.nlm.nih.gov/sra/?term=PRJNA434002, https://cellxgene.cziscience.com/collections/d5d0df8f-4eee-49d8-a221a288f50a1590, https://cellxgene.cziscience.com/collections/283d65eb-dd53-496d-adb7-7570c7caa443.

The raw data are also available in the respective original publications.

#### Data Compatibility

To preserve comparability and reproducibility, we used each dataset as released by the original authors and applied additional quality control, preprocessing, and re-normalization steps to ensure consistency across studies. The integrated dataset was constructed using only the genes common to all individual datasets (13,746 features).

Notably, PPEF1 was consistently detected in 17 of the 18 datasets (in-house and publicly-available), demonstrating its robust expression across studies and preprocessing pipelines. In the integrated object, the raw.X layer contains 13,747 features (including PPEF1) with imputed zero values in the Prashant et. al. dataset, which had excluded PPEF1 during its original preprocessing. Because these counts are imputed and experimentally measured counts are not available, PPEF1 values from the Prashant dataset should not be used for quantitative downstream analyses. When the Prashant dataset is excluded, PPEF1 expression values from the remaining datasets can be used for downstream processing. Figures 2H–2I were generated after excluding the Prashant et. al. dataset.

#### Data Processing and Quality Control

All analyses were conducted using Python 3.9 (Scanpy v1.10.3) and R 4.4 (Seurat v5.2.1). Metadata were curated manually to ensure consistency across datasets.

Raw counts were imported into Scanpy as AnnData objects. Per-cell quality control metrics were computed, including the total number of detected genes (n_genes_by_counts) and the percentage of mitochondrial transcripts (percent_mt). Mitochondrial genes were identified by the prefix “MT-”.

Cells were filtered using the following criteria:

1. Fewer than mean − 1.2 × standard deviation detected genes were excluded.
2. Cells with > 5% mitochondrial content were excluded.

Over 90% of cells per sample typically passed these filters. Datasets that had undergone prior filtering were reviewed but not reprocessed. Cells with mitochondrial content >5% were consistently removed.

After filtering, raw counts were normalized to 10,000 counts per cell, log-transformed, and highly variable genes (HVGs) were identified using the Seurat method (n_top_genes = 5000). Expression values were scaled to a maximum of 10 prior to PCA (50 components). Nearest neighbors (20–40) and UMAP embeddings were computed, and clusters were identified using the Leiden algorithm across multiple resolutions.

Harmony batch correction (harmonypy v0.1.0) was performed using sample_id as the batch variable. Post-harmonization, QC metrics were re-evaluated and confirmed to be consistent across datasets. To quantitatively assess batch mixing after integration, we computed the rejection rates using the UMAP embedding, with the scib package. Lower rejection rates indicate better local mixing of cells from different batches, and the observed values confirmed that harmony effectively removed batch-specific structure.

#### Integration and Downstream Analysis

Ensembl gene identifiers were converted to gene symbols using MyGene.info; genes without mappings were excluded. Individual datasets were concatenated into a merged AnnData object using outer joins to preserve shared genes. Normalization, HVG selection, and scaling were repeated on the merged data. PCA was performed (15–50 components), and Harmony correction was applied to remove batch effects. The Harmony-corrected embedding (X_pca_harmony) was used for neighbor graph construction (20 neighbors) and UMAP visualization.

Clusters were identified using Leiden clustering (multiple resolutions). Metadata in .obs were curated to include sample_id, dataset, organism, region, subregion, matter (white or gray), tissue, condition, and sex. Low-quality clusters were manually reviewed and excluded. UMAP, dot plots, and violin plots were generated using matplotlib and seaborn (v0.13) to visualize marker expression and QC metrics.

##### Cell Type and Basic Cluster Annotation

Differential gene expression analysis was performed using the Wilcoxon rank-sum test (scanpy.tl.rank_genes_groups) on log-normalized data. For each cluster, test statistics including log fold-change, adjusted p-value, and standardized test score were extracted.

Marker genes were ranked based on Wilcoxon “scores”, which capture both the magnitude (log₂ fold change) and consistency of expression differences between the target cluster and all others. Higher scores indicate stronger and more reliable enrichment within that cluster.

Clusters were annotated to canonical cell types based on established marker genes:

- <u>Neurons:</u> *SYT1, SNAP25*
- <u>Oligodendrocytes:</u> *MBP, ST18*
- <u>OPCs:</u> *PDGFRA, PCDH15, VCAN*
- <u>Schwann cells:</u> *MPZ, PMP22*
- <u>Microglia:</u> *CD74, CX3CR1, LRMDA, ARHGAP15*
- <u>Macrophages:</u> *F13A1, CD163, LRMDA, ARHGAP15*
- <u>Immune cells:</u> *STAT4, THEMIS, ARHGAP15*
- <u>Astrocytes:</u> *SLC1A2, GFAP, AQP4*
- <u>Ependymal cells:</u> *FOXJ1, DYNLRB2, DCDC1*
- <u>Endothelial cells:</u> *VWF, FLT1, CLDN5*
- <u>Mural cells:</u> *PDGFRB, RGS5*
- <u>Fibroblasts:</u> *CEMIP, DCN, LAMA2*

Cell-type annotations were validated by comparing the marker-based labels with results from post hoc subclustering and manual review, showing >95% concordance across methods.

#### Annotation of Activation States

To assess oligodendrocyte activation, subsets of cells from the spinal cord and motor cortex were reanalyzed independently. Expression values were normalized, log-transformed, and scaled as described above.

A predefined gene panel (*CD9, FTH1, CNP, CRYAB, FTL, MAG, MARCKSL1, NKX6-2, RNASE1, APOD*) was used to compute per-cell activation scores. A Gaussian Mixture Model (GMM) with two components was fitted to the activation score distribution using scikit-learn (n_components=2, covariance_type=’full’, n_init=10) per condition. The higher-expression component was designated “reactive,” and the lower component “homeostatic.” The trained GMM was then applied to all samples to assign Activation_State labels.

These labels were validated via independent subclustering and marker-based post hoc annotation, confirming strong agreement across methods. UMAPs were used to visualize the spatial segregation of activation states.

#### Manual Cluster Curation and Doublet Inspection

Clusters expressing multiple cell-type markers were manually inspected using UMAP feature and dot plots. Doublet detection was initially performed using DoubletFinder and scDblFinder, but manual curation proved more reliable. In large datasets, low-quality or doublet-like cells formed distinct clusters that were easily detected through visual inspection.

#### Functional Enrichment Analysis (Figure 1–2)

Differentially expressed genes (DEGs) were ranked by Wilcoxon score, and the top 10% were used for enrichment testing. Gene symbols were mapped to Entrez IDs using clusterProfiler (v4.10.1) and org.Hs.eg.db (v3.19.1).

Enrichment analyses included:

- <u>Gene Ontology (GO):</u> enrichGO and groupGO for biological processes.
- <u>KEGG:</u> enrichKEGG and gseKEGG.
- <u>Reactome:</u> enrichPathway and gsePathway.
- <u>Disease Ontology (DO):</u> enrichDO.

Redundant GO terms were reduced (cutoff = 0.8). Visualizations included lollipop plots, Venn diagrams (*eulerr*), and UpSet plots (*ComplexUpset*). All enrichment analyses were performed in R 4.4 using clusterProfiler, ReactomePA, DOSE, and ggplot2.

#### Statistical Analysis for Group Comparisons (Figure 1–2)

Per-sample proportions of oligodendrocyte subpopulations were calculated separately for each region.

Distributions were assessed for normality using the Shapiro–Wilk test (scipy.stats). Given the small sample size and non-normal distributions (p < 0.05), all group comparisons were performed using the nonparametric Mann–Whitney U test (scipy.stats).

Multiple testing correction was applied using the Benjamini–Hochberg false discovery rate (FDR) procedure (statsmodels).Significance thresholds were defined as: * p.adj < 0.05; ** p.adj < 0.01; *** p.adj < 0.001.

#### Enzyme-Linked Immunosorbent Assay (ELISA) (Figure 3A–3B)

ELISAs were performed to quantify LRP2, CNP, MBP, NFL, and pNFH in cerebrospinal fluid (CSF).

- <u>Sample sizes:</u> LRP2 (control n=18; sALS n=18; C9ALS n=8), CNP (control n=17; sALS n=10; C9ALS n=10), MBP (control n=17; sALS n=10; C9ALS n=8), NFL (sALS n=25; C9ALS n=9), and pNFH (sALS n=25; C9ALS n=9).
- Measurements were performed in duplicate, and results differing by >1 SD between replicates were excluded.
- Commercial kits were used following manufacturer’s protocols:

- <u>LRP2:</u> BIOMATIK (EKF58729)
- <u>CNP:</u> assaygenie (HUFI01436)
- <u>MBP:</u> Ansh Labs (AL-108)
- <u>NFL:</u> Uman Diagnostics (10-7001)
- <u>pNFH:</u> EUROIMMUN (EQ 6561-9601)

Absorbance was measured at 450 nm, and concentrations were interpolated from standard curves.

#### Quantification and Visualization of ELISA Data (Figure 3A–3B)

ELISA results were processed in Python 3.9 (pandas v2.2, numpy v1.26). Missing data were coded as “Missing.”

Given the small group sizes and non-normal distributions, Kruskal–Wallis tests were used for group comparisons, followed by pairwise Mann–Whitney U tests with Benjamini–Hochberg FDR correction. Correlations between markers were assessed using the Spearman rank correlation coefficient (ρ) within each group.

Plots were generated using seaborn v0.13 and matplotlib v3.8, displaying combined box–strip plots with annotated significance levels.

#### Compositional Analysis (Figure 3C–3R)^115^

Given the inherent complexity and statistical challenges of compositional analyses in single-cell datasets, we employed a multi-model strategy to identify and validate differential abundance (DA) patterns across cell populations. Two main categories of compositional frameworks were used.

**The first category** comprised cluster-based methods, which rely on predefined cell-type or cluster annotations. These included:

<u>scCODA</u>, a Bayesian hierarchical model for compositional inference, was applied to quantify shifts in cellular composition across diagnostic contrasts. For each region (motor cortex and spinal cord) and condition (Control vs. sALS or C9ALS), nuclei counts were aggregated by sample and cell type to construct sample-by-cell-type matrices. Differential abundance was estimated using scCODA (v0.1.9) via Hamiltonian Monte Carlo sampling (20,000 iterations), testing all possible reference cell types to identify consistently credible compositional changes while accounting for the inherent compositional structure of single-cell data.

<u>DCATS</u>, a beta-binomial regression framework, was used as an independent method to assess compositional variability while explicitly modeling overdispersion and sample-level heterogeneity. For each region and diagnostic comparison (Control vs. sALS; Control vs. C9ALS), sample-by-cell-type count matrices were analyzed using DCATS, incorporating a cluster-level similarity matrix derived from the Seurat k-nearest neighbor graph where available. Significance was assessed according to package defaults, and results were cross-validated against scCODA and propeller outputs to ensure robustness.

<u>Propeller</u>, an empirical Bayes method implemented in the speckle package, was further used to test for differences in cell-type proportions across groups. Single-nucleus counts were aggregated into sample-by-cell-type tables for each contrast (Control vs. sALS; Control vs. C9ALS). Analyses were performed using both logit and arcsine–square-root transformations to stabilize variance, with robust variance estimation applied. Moderated t-tests or ANOVA were conducted following propeller defaults, and resulting statistics were compared across compositional frameworks (scCODA, DCATS) for validation.

Finally, <u>scDC</u>, a bootstrap-based compositional modeling framework, was employed to quantify uncertainty and test for differential cell-type abundance. Normalized single-nucleus expression data were analyzed using scDC (v1.0.1) with the scDC_noClustering function, performing 200 bootstrap resamples and computing percentile, BCa, and multinomial confidence intervals for each cell type. Subject-level proportions were compared between diagnostic categories (Control vs. sALS; Control vs. C9ALS) using generalized linear models implemented in scDC::fitGLM. Multiple testing correction was applied using the Benjamini–Hochberg procedure (FDR < 0.05). Complementary analyses were performed using scDC_clustering, which incorporates transcriptional similarity between cell types into the compositional model for increased robustness.

**The second category** included cluster-free approaches that assess DA at the neighborhood level without relying on predefined clusters. These methods—Milo, DA-seq, and MELD—detect spatially localized changes in cell abundance directly from the embedding space.

<u>Milo</u>, a neighborhood-level differential abundance framework, was used to identify spatially localized compositional changes independent of predefined clusters. Single-nucleus expression data were converted to SingleCellExperiment objects and analyzed using MiloR (v1.9.1). A k-nearest neighbor graph was constructed from the PCA embedding (k = 30, 40 dimensions), and overlapping cellular neighborhoods were generated (prop = 0.05). For each sample, neighborhood counts were aggregated to form a sample-by-neighborhood matrix. Differential abundance testing was performed using the testNhoods function with a design formula (∼ Status), incorporating sample-level metadata. Neighborhoods with a spatial false discovery rate (SpatialFDR) < 0.05 were considered significantly altered. Significant neighborhoods were annotated by cell class, and neighborhood-level DA patterns were visualized using UMAP overlays and beeswarm plots. Marker genes distinguishing DA neighborhoods were further identified using the findNhoodGroupMarkers function on log-normalized expression values.

##### Integration and cross-validation

To improve robustness and reduce false-positive findings (Type I errors), only compositional results supported by at least two of the three retained cluster-based methods (scCODA, DCATS, propeller) were considered significant. Among these, DCATS and propeller demonstrated higher sensitivity—particularly for underrepresented populations—whereas scCODA adopted a more conservative modeling strategy. Accordingly, the primary figures display results derived from scCODA, while all reported findings were validated by at least two independent frameworks. scDC was excluded from downstream interpretation due to a high false-positive rate.

For neighborhood-level analyses, results from Milo were cross-compared with DA-seq and MELD, which exhibited high consistency in most datasets. However, because DA-seq and MELD performance can decline in the presence of batch effects, Milo results were prioritized for final interpretation.

##### Quality control and visualization

To avoid confounding by experimental enrichment or biased sampling, datasets annotated as having enrichment (“Enrichment” == “Yes”) were excluded from all compositional analyses.

For visualization and independent validation, compositional analysis of cell-type abundance was performed to quantify the relative proportion of oligodendrocytes across diagnostic groups (Control, sALS, C9ALS) in the spinal cord. Cell-level annotations were derived from the harmonized AnnData object. Only non-enriched spinal cord samples were retained; “fALS” and “sALS_FTD” samples were excluded.

For each sample, the total number of nuclei and the number annotated as Oligodendrocytes were computed from the .obs metadata. Per-sample proportions were calculated as: Proportion_oligo = n_oligo / n_total

This yielded one compositional value per biological replicate. Proportions were then joined to corresponding sample-level metadata to retain diagnostic group information.

All statistical analyses and visualizations were conducted in Python 3.9 using scanpy v1.10.3, pandas v2.2, numpy v1.26, and seaborn v0.13. Box–strip plots were generated to depict per-sample proportions across diagnostic groups, with each dot representing an individual sample and boxplots summarizing interquartile ranges. Outliers were suppressed for clarity, and a consistent color palette was applied across all groups.

#### Pseudobulk Differential Gene Expression Analysis (Figure 4–S2)

Pseudobulk differential gene expression (DGE) analysis was performed to assess transcriptional alterations in oligodendrocytes within the spinal cord. Analyses were restricted to non-enriched (Enrichment == “No”) samples annotated as control, sALS, or C9ALS. Genes expressed in ≥20% of nuclei in at least one diagnostic group were retained. To reduce noise from lowly expressed genes, the bottom 20th percentile by mean expression was excluded, retaining the top 80% of expressed features.

For each biological replicate, single-nucleus counts were aggregated by Sample_ID to generate a pseudobulk expression matrix. Corresponding metadata were joined to preserve sample-level annotations, including Sex, Dataset, and Status. Differential expression testing was conducted using PyDESeq2 (v0.3.1), a Python implementation of DESeq2, with the design formula ∼ Dataset + Sex + Status. Dispersion estimates and size factors were computed using the Wald test. Pairwise contrasts were performed between sALS and Control samples, and similarly for other diagnostic comparisons.

Significance was defined as Benjamini–Hochberg adjusted p (FDR) < 0.05 and absolute log₂ fold-change > 0.25. Differentially expressed genes were visualized using volcano plots, dot plots, and heatmaps.

Marker gene expression patterns were further visualized using Scanpy (v1.10.3). Normalized and batch-corrected expression values were obtained from the scvi_expr layer generated by scVI. Samples were ordered by diagnostic category (C9ALS, control, sALS). For each gene, color intensity indicates the average scaled expression level, and dot size represents the fraction of nuclei expressing that gene within each sample.

Matrix plots were generated to visualize mean expression levels of canonical marker genes across samples and diagnostic groups. Expression values were obtained from the log1p_normalized layer of the harmonized AnnData object, corresponding to log-transformed and normalized counts. Samples were ordered by diagnostic category (C9ALS, control, sALS) to ensure consistent visualization across datasets. For each gene, color intensity represents the average scaled expression across nuclei within each sample. Marker genes were selected to capture canonical lineage and activation signatures.

Plots were generated using the sc.pl.matrixplot function in Scanpy (v1.10.3) with standardized scaling across variables (standard_scale=“var”) and the continuous Spectral_r colormap. Aesthetic adjustments were applied in Matplotlib (v3.8) to harmonize figure style, remove spines and gridlines, and standardize font formatting. Figures were exported as high-resolution, transparent PNG files (800 dpi) for inclusion in final figure panels.

Gene Ontology (GO) enrichment analysis was performed as described in the Functional Enrichment Analysis section. Genes meeting the thresholds of padj < 0.05 and |log₂ fold-change| > 0.3 were used as input. GO terms associated with biological processes were identified using clusterProfiler and summarized to highlight pathways most enriched among the upregulated and downregulated gene sets.

#### Trajectory Inference. (Figure 5)

Trajectory and lineage inference analyses were conducted to characterize the transitions between homeostatic and reactive oligodendrocyte states and to ensure robust directional inference of pseudotemporal progression. Multiple complementary frameworks were applied to assess trajectory directionality and to validate lineage relationships.

##### Initial trajectory inference

Monocle3 (v1.3.1) and Slingshot (v2.10.0) were used to infer primary lineage structures. Monocle3 constructs principal graphs in UMAP space, while Slingshot fits smooth principal curves in PCA space. Cluster labels corresponding to OPCs, oligodendrocytes, and subclusters were used as anchors for lineage assignment. Both methods produced consistent pseudotime orderings from OPC-like to mature oligodendrocyte states.

###### Monocle3

Integrated Seurat objects were converted into Monocle3 CellDataSet objects using the as.cell_data_set function. Cells were clustered (cluster_cells), and principal graphs were learned within the UMAP embedding using learn_graph, with a minimum branch length of 10 to prevent overfitting. Root principal nodes were manually defined based on progenitor populations, and pseudotime values were computed using order_cells. Pseudotime trajectories were visualized with plot_cells, and mean pseudotime values were summarized per cluster to capture progression. For three-dimensional visualization, UMAP embeddings were reconstructed using Seurat v5.0.0 (RunUMAP, 40 PCs, Harmony reduction, n.components = 3) and plotted with plot_cells_3d.

###### Slingshot

Seurat objects were converted to SingleCellExperiment objects containing UMAP embeddings. Lineages were initialized from the “RG” or “Olig_1” clusters, and pseudotime was inferred along smooth principal curves. Cluster-level mean pseudotime values were computed and visualized as gradients across the UMAP embedding, with fitted trajectory curves overlaid.

##### Condition-associated differences in trajectory dynamics

###### Lamian

Integrated Seurat objects were converted for Lamian analysis using PCA embeddings and log-normalized expression values. Trajectory structures were inferred using infer_tree_structure, with PDGFRA-defined OPC-like cells set as the root population. Uncertainty was assessed via permutation tests (n = 100–1,000). Condition effects were evaluated using branchPropTest and lamian_test, and significant trajectory–condition interactions were visualized using getPopulationFit, clusterGene, and plotXDEHm.

##### Diffusion-based pseudotime and transcriptional entropy

###### Palantir

Analyses were conducted on log-normalized matrices following MAGIC imputation. Pseudotime and entropy were computed using run_palantir (knn = 30), and cluster-wise mean values were summarized to capture global activation dynamics.

##### Probabilistic latent-time modeling and directed graph reconstruction

###### MARGARET

The model was trained on MAGIC-imputed matrices using 30 episodes and 10 metric-learning epochs. Directed transition graphs were generated from pseudotime-ranked cluster connectivity matrices, and transition probabilities were visualized using Scanpy and Matplotlib.

##### Graph topology validation

###### Partition-based Graph Abstraction (PAGA)

Harmony-reduced embeddings (X_harmony) were used to compute kNN graphs (n_neighbors = 30, n_pcs = 10). PAGA-derived connectivity structures were consistent with MARGARET-inferred trajectories. Hierarchical lineage relationships were visualized using radial tree layouts.

##### Robustness and statistical assessment

Directionality and transition robustness were quantified using two complementary strategies. First, per-donor pseudotime-oriented transition flows were recomputed independently, and homeostatic → reactive versus reactive → homeostatic transition magnitudes were compared using linear mixed-effects models (donor as random effect). Second, bootstrap resampling (1,000 iterations) was applied to estimate 95% confidence intervals for the net directional flow difference (H→R − R→H). Directional trends were retained only when supported by both donor-level and bootstrap-based analyses.

##### Cross-validation and tissue-specific analyses

All trajectory analyses were conducted on a merged dataset combining motor cortex and spinal cord oligodendrocytes to increase sample size and model stability. Results were validated separately in each tissue to ensure consistency. For ALS datasets, C9ALS and sALS spinal cord samples displayed similar dynamics and were merged for trajectory inference, whereas C9ALS and sALS motor cortex samples exhibited greater variability and were analyzed independently. Across all trajectory inference methods—including Monocle3, Slingshot, Palantir, MARGARET, Lamian, and PAGA—the pseudotemporal orderings were largely consistent. This convergence supports a clear directional progression from homeostatic to reactive oligodendrocyte states in control samples, and the reverse in the disease spinal cord. Notably, this agreement is critical for MARGARET, as it relies on the pseudotime-derived directional information to construct its trajectory arrows.

#### RNA Velocity, Fate Mapping and Dynamics. (*Figure 5*–S3)

##### RNA velocity data preprocessing and integration

Raw lane-level .loom files containing spliced and unspliced count matrices were imported into AnnData objects using Scanpy. Each lane was annotated with its corresponding barcode and curated sample metadata. Cells were subset to retain only those present in the prefiltered reference object, and duplicated gene names were removed prior to merging. Lane-level datasets were concatenated and subset to features common with the reference dataset, then preprocessed and saved for downstream RNA velocity inference.

##### RNA velocity Inference

RNA velocity analyses were performed using scVelo (v0.2.6). The preprocessed and harmonized AnnData object containing spliced and unspliced counts was used as input. Gene-wise dynamical models were estimated with scv.tl.recover_dynamics, and velocities were computed using scv.tl.velocity with the dynamical model. Velocity graphs and embeddings were visualized on UMAP, with confidence filtering applied to retain cells with high-confidence velocity vectors (velocity_confidence > 0.6). Latent time and gene-wise dynamics were inferred (scv.tl.latent_time, scv.tl.rank_dynamical_genes), and gene-specific velocity profiles were plotted for selected genes of interest. Pseudotime was computed using scv.tl.velocity_pseudotime, and PAGA-based connectivity was evaluated to infer cluster-level transitions. Kinetic parameters (transcription, splicing, degradation rates) were extracted and summarized, and differential kinetic testing was performed across clusters to identify genes with cluster-specific dynamics. Velocity magnitudes and confidence metrics were visualized across clusters and regions, and summary statistics were exported for downstream analyses.

##### CellRank Analysis and Fate Probabilities

To infer cell-state transitions and fate probabilities, RNA velocity–derived transition matrices were computed using the VelocityKernel from CellRank (v1.7.0) and integrated with connectivity information via the ConnectivityKernel. Pseudotime was estimated along the inferred velocity graph, and root cells were manually defined based on annotated homeostatic oligodendrocytes. Initial and terminal states were predicted using GPCCA, and macrostates were visualized on UMAP and PHATE embeddings. Differential expression along terminal versus non-terminal states was assessed using violin plots and t-tests. Fate probabilities were computed for terminal macrostates, and circular projections were used to visualize lineage relationships across clusters and subclusters.

##### RNA Velocity and Vector Field Reconstruction

To further characterize transcriptional dynamics and cell state transitions, we applied Dynamo (v1.3.2) to the single-cell dataset. Unspliced RNA counts were used as the primary expression matrix, and data were preprocessed following a Monocle-based workflow. The top 5,000 highly variable genes were selected for downstream analyses, and cell cycle scoring was disabled. We re-run the analysis 5 times on different batches of high variable genes, as well as to the whole dataset.

Gene-wise dynamics were modeled using the dynamical framework in Dynamo, estimating transcription, splicing, and degradation rates per gene. Cell-wise velocity vectors were computed using Pearson correlation between observed and modeled gene expression, and confidence scores were assigned based on lineage-specific dynamics.

Velocity vectors were projected onto UMAP embeddings, and visualized using scatterplots, streamlines, and phase portraits for top dynamical genes. Streamline and quiver plots highlighted directional flows across cell populations, with arrow sizes and lengths adjusted for clarity. Confidence filtering was applied to retain robust cell-wise velocities.

A vector field was reconstructed for PCA and UMAP embeddings to characterize local and global cell state transitions. Metrics including speed, curl, divergence, acceleration, and curvature were computed to capture dynamic properties of the system. Energy landscapes and DD-Hodge potentials were calculated to estimate the directional potential governing cell state transitions. Cell-cluster-specific distributions of DD-Hodge potential were summarized with boxplots, and statistical differences were assessed using mixed-effects linear models with sample identity as a random effect.

##### Cell-level Entropy Quantification

To quantify transcriptional disorder and infer cell-state stability, we applied the scEntropy framework on normalized single-cell transcriptomes. The log-transformed expression matrix (log1p_normalized layer) from the merged AnnData object was used as input. For each cell, scEntropy was computed using the RCSA (Reference Cell State Average) option, which evaluates the divergence of each cell’s expression profile from the system’s mean reference vector, thereby providing a global measure of transcriptional disorder.

Entropy distributions were compared across cellular subtypes and activation states. Group-level differences in entropy were assessed using linear mixed-effects models with sample identity as a random effect (statsmodels.mixedlm), controlling for inter-sample variability. Gene-level entropy was additionally computed using histogram-based Shannon entropy on binned expression values, identifying highly variable and regulatory genes contributing to system-level instability. Visualization of entropy distributions across oligodendrocyte subpopulations and regional clusters was performed using Seaborn boxplots and jittered scatter overlays.

#### Gene Regulatory Network Analysis. (*Figure 6*)

##### Network Construction

###### CellOracle

To reconstruct oligodendrocyte gene regulatory networks (GRNs), filtered single-nucleus RNA-seq data were imported into CellOracle (v0.10.12), retaining only genes present in the promoter-based human transcription factor (TF) network. Highly variable genes and TF-target relationships were used to construct cluster-specific regulatory matrices. Principal component analysis (PCA) was performed on the raw count matrix using truncated singular value decomposition (SVD), followed by k-nearest neighbor (KNN) imputation to smooth gene expression and account for sparsity in large datasets

Cluster-specific GRNs were inferred using CellOracle’s get_links function with significance thresholds (p < 0.001, top 2,000 weighted links), and network scores—including degree, eigenvector centrality, and betweenness—were computed. Filtered link tables were exported for each cluster for downstream analysis and visualization.

Because CellOracle is optimized for smaller datasets (1,000–3,000 cells and a few thousand genes), we first ran the analysis on the full oligodendrocyte dataset and then repeated it 20 times using bootstrap resampling, with batches of 3,000 cells and 3,000 genes, which yielded consistent results.

###### SCENIC

We next applied SCENIC (v0.12.1)) to independently reconstruct gene regulatory networks and validate key transcriptional drivers identified with CellOracle. Whereas CellOracle estimates transition-associated regulatory influences by integrating TF–promoter priors with expression dynamics, SCENIC infers regulons—TFs and their direct target genes—based on co-expression patterns followed by cis-regulatory motif enrichment, providing a complementary and orthogonal assessment of TF activity.

To reduce noise and computational burden, we first quantified expression across all annotated human transcription factors and retained the top 25% most highly expressed TFs in the dataset. Using these TFs, we downsampled the full dataset to ∼100,000 cells while preserving sample and cluster composition, and extracted the top 5,000 highly variable genes (plus all retained TFs) for GRN inference. Expression matrices were normalized, log-transformed, and exported for GRNBoost2 (SCENIC Step 1), which was executed to generate TF–target adjacency lists.

Motif enrichment (SCENIC Steps 2–3) was performed on the full dataset, using cis-regulatory ranking databases (10 kb upstream/downstream and promoter-proximal windows) together with motif annotations. Enriched motifs were filtered based on normalized enrichment scores (NES ≥ 2.0) and direct or orthology-supported motif–gene annotations. Regulons were derived from high-confidence TF–target modules (≥5 genes) and renamed according to their corresponding TFs.

To account for potential stochasticity in GRNBoost2 and the motif enrichment procedure, the full SCENIC workflow (Steps 1–4) was repeated 20 independent times across both full and subsampled datasets, in both only control samples and the whole dataset (control and disease). Regulons consistently recovered across iterations were retained for downstream analyses, ensuring robustness to random seed variation and sampling noise.

Regulon activity was quantified for each cell using AUCell, and both continuous activity scores and binarized activity states were computed. The resulting AUC matrix was Z-transformed to enable comparison across regulons. To identify regulators driving differences between activation states, we fitted linear mixed-effects models for each regulon, modeling regulon activity as a function of cluster identity while treating sample ID as a random effect. Multiple testing correction was performed using the Benjamini–Hochberg method. Regulons showing significant differential activity across states were cross-referenced with TF differential expression results and CellOracle-inferred drivers to identify high-confidence, convergent regulators of oligodendrocyte activation trajectories.

This SCENIC workflow, implemented across full and subsampled datasets, provided an independent validation layer for regulatory programs identified in CellOracle and increased confidence in TFs underlying state transitions.

##### Differential Network Analysis (DNA)

To compare regulatory architecture across biological conditions, we performed differential network analysis using CellOracle-derived TF–target edges from control motor cortex, control spinal cord, C9ALS, and sALS datasets. For each condition, regulatory edges were represented as weighted TF–target pairs using the CellOracle coef_mean statistic. To ensure comparability across networks, we first generated a unified gene list comprising the union of all TFs and target genes identified in any condition. All initial CellOracle networks were then filtered to retain only edges whose source and target nodes were present in this unified gene set, ensuring consistent node universes across control and disease networks.

Networks were reconstructed using networkx, with edges weighted by regulatory coefficients. To focus on biologically meaningful regulatory changes, we restricted the feature space to TFs expressed in at least 25% of cells within each oligodendrocyte subcluster and to target genes that were both expressed in ≥10% of cells and differentially expressed between activation states. Although this filtering may omit regulatory interactions that are not transcriptionally dynamic between states, repeating the analysis using the full unfiltered TF–target space confirmed that key TFs—such as JUND—remained consistently prioritized.

###### Edge-Level Differential Analysis

For each pairwise comparison (ALS vs. control), we standardized TF–target interactions by defining an undirected edge identifier for every link. Edge tables were merged across conditions, and differential network metrics were computed: Δ coefficient, Δ –logp, Z-scored Δ coefficient, standardized across all edges

Edges with |Δ coefficient| > 0.1 and |Δ –logp| > 1 were considered significantly altered, representing either gained or lost regulatory interactions. These edges were visualized using a Kamada–Kawai force-directed layout, with edge width proportional to |Δ coefficient| and edge color reflecting directionality of change.

###### Node-Level Differential Analysis

To identify regulators with altered connectivity in ALS, we computed multiple centrality metrics for each node (TF or target gene): degree centrality, betweenness centrality, eigenvector centrality (when supported by graph structure)

Node-level differential metrics (e.g., Δ degree) were used to rank genes based on changes in their regulatory importance. Analyses were performed both on all nodes and specifically on TFs (“source nodes”), since TFs represent direct regulators of transcriptional programs.

###### Network Topology and Community Structure

Global network properties—including node count, edge count, density, and transitivity—were computed for each condition. Community structure was assessed using the greedy modularity optimization algorithm. Differences in topology between ALS and control networks were statistically evaluated using:

– Wilcoxon rank-sum test for density and transitivity
– Kolmogorov–Smirnov test for degree distribution shifts

This allowed quantification of both local (edge-level) and global (network-level) alterations in regulatory architecture associated with ALS.

###### Monte Carlo Significance Testing

To evaluate whether observed differences in degree or betweenness centrality exceeded those expected by chance, we implemented a Monte Carlo network rewiring procedure. For each condition pair, networks were independently randomized 1,000 times using degree-preserving double-edge swaps (20% of edges per iteration). For each randomized network pair, mean degree and betweenness centrality were recomputed, generating empirical null distributions. Observed centrality differences were then compared to these null distributions to derive empirical p-values, providing a robust statistical framework for assessing ALS-specific changes in network topology.

###### Visualization

Network visualizations were generated using networkx, matplotlib, and seaborn in Python, complemented by igraph, tidygraph, and ggraph in R for higher-order graph layouts. Differential centrality plots, jittered barplots, and heatmaps were produced using custom plotting functions.

#### Metabolic Pathway Activity Analysis. (Figure 7)

Metabolic pathway activity was quantified using scMetabolism (v0.2.1), applying KEGG and Reactome pathway annotations to the processed Seurat object. To maintain compatibility across workflows, the RNA assay was duplicated before scoring. Pathway activity was computed using the AUCell framework, which ranks genes by expression within each cell and quantifies the enrichment of pathway-associated genes within the high-expression tail.

For both KEGG and Reactome databases, AUCell scores were stored in a dedicated METABOLISM assay and visualized across clusters, activation states, and disease groups using UMAP overlays, dot plots, and boxplots. Per-cell pathway activity values were reshaped into long format to enable flexible pathway-level visualization and phenotype-specific normalization.

To identify metabolic pathways differentially active across oligodendrocyte states or disease conditions, we used linear mixed-effects models, treating pathway activity as the response variable, the phenotype of interest (e.g., activation state, status) as the fixed effect, and Sample_ID as a random intercept to account for donor-level variability and avoid pseudoreplication. P-values for fixed effects were adjusted using the Benjamini–Hochberg method.

#### Cell-cell Communication Analysis. (Figure 7)

##### CellChat

To infer intercellular signaling, we applied CellChat (v2.1.2) on our Seurat object, grouping cells by cluster. We initialized a CellChat object and set the ligand–receptor database to CellChatDB.human. Over-expressed ligands and receptors across groups were identified using identifyOverExpressedGenes and identifyOverExpressedInteractions, respectively, following the recommended preprocessing outlined in the CellChat tutorial.

We then computed communication probability using a protein-protein interaction (PPI) graph (smoothData), filtered low-confidence interactions (minimum cells = 10), and inferred pathway-level communication with computeCommunProbPathway. Networks were aggregated (aggregateNet) to summarize both the number of interactions and interaction strength between cell groups.

For visualization, we used circle plots to display overall communication networks (netVisual_circle), and hierarchical and chord diagrams (netVisual_aggregate) for specific signaling pathways (e.g., “PSAP”). We also assessed cell-group signaling roles by computing centrality metrics on pathway-level networks (netAnalysis_computeCentrality) and visualizing them via scatter plots or heatmaps (netAnalysis_signalingRole_scatter and netAnalysis_signalingRole_heatmap).

To investigate individual ligand–receptor contributions, we extracted enriched LR pairs (extractEnrichedLR) and visualized them using chord, bubble, and individual network layouts.

##### LIANA+

To investigate intercellular communication among oligodendrocyte subpopulations, we performed ligand–receptor inference using the LIANA (v0.1.14) framework. The full single-cell RNA-seq dataset was first converted into a SingleCellExperiment object. LIANA was executed using multiple inference methods—including Cytotalk, Connectome, LogFC, NATMI, and SingleCellAggregator (SCA)—with the consensus ligand–receptor database and a minimum expression threshold of 10 cells per cluster. For each method, ligand and receptor expression proportions were calculated, and 100 permutations were used to assess interaction significance. Individual method results were aggregated into a consensus score using LIANA’s rank aggregation procedure. To focus on high-confidence interactions, we retained ligand–receptor pairs with a magnitude rank ≤ 0.05. Interaction networks were visualized using dot plots, density distributions of method scores, heatmaps of selected ligand–receptor pairs (e.g., SPP1), and chord diagrams representing cluster-to-cluster communication.

For orthogonal validation and downstream analysis of signaling pathways, we integrated LIANA results with NicheNet. Expression matrices and cluster metadata were used to define potential sender (ligand-producing) and receiver (target) populations. The ligand–target regulatory matrix was retrieved from the published NicheNet database, and potential ligands were defined based on prior knowledge and LIANA-inferred interactions. The top differentially expressed genes in receiver populations were identified using standard thresholds (adjusted p < 0.05, pct.1 > 0.2) and used as input target gene sets for ligand activity prediction. Predicted ligand activities were quantified using Pearson correlation scores between ligand expression and target gene response, and visualized alongside receptor engagement using heatmaps and bar plots.

NicheNet was also run independently of LIANA to directly predict ligand activity for specific target gene sets, providing an additional validation layer for inferred ligand–receptor interactions and ensuring robust identification of key signaling pathways.

All visualizations, including dot plots, heatmaps, chord diagrams, and receptor–ligand activity plots, were generated using the ggplot2, Complex Heatmap, and cow plot R packages.

##### NicheNet

To complement CellChat and LIANA-inferred ligand–receptor interactions, we performed independent NicheNet analyses to prioritize upstream ligands and their putative target genes in receiver populations. Single-cell RNA-seq data were imported into NicheNet, and expressed genes were defined per receiver cluster (pct ≥ 0.5) and per sender cluster (pct ≥ 0.1). Potential ligands were filtered based on expression in senders and inclusion in the NicheNet ligand–target matrix. Differentially expressed genes in receiver clusters were used as the gene set of interest, with background genes defined as expressed genes overlapping the ligand–target matrix.

Ligand activities were quantified using predict_ligand_activities, generating AUPR-ranked ligands. Top-ranked ligands were visualized via heatmaps and integrated with predicted target genes using get_weighted_ligand_target_links. Receptor–ligand interactions were visualized with get_weighted_ligand_receptor_links. For sender-focused analyses, potential ligands were further restricted to those expressed in specific sender cell populations. Heatmaps and dot plots were used to visualize ligand activities, target gene regulation, and receptor interactions.

#### Statistical Testing for Figures 5 to 7. (Figure 5-7)

All statistical comparisons presented in Figures 5, 6 and 7—evaluating differences in metrics such as entropy, pseudotime, regulon activity, and metabolic scores—were performed using generalized linear mixed-effects models (GLMMs). Analyses were conducted in Python using statsmodels.formula.api, with donor identity included as a random effect to account for inter-donor variability. For comparisons involving multiple groups, such as metabolic scores, p-values were adjusted using the Benjamini–Hochberg method.

## ACKNOWLEDGMENTS

This work was supported by grants from KU Leuven (C1 - C14/22/132), Opening the Future Fund (KU Leuven), the ALS Liga België and Target ALS (BM-2024-C3-L1).

PVD holds a fundamental clinical investigatorship of KU Leuven and is supported by the E. von Behring Chair for Neuromuscular and Neurodegenerative Disorders and the KU Leuven funds “Een Hart voor ALS” and “Laeversfonds voor ALS Onderzoek”. DRT received funding from Fonds Wetenschappelijk Onderzoek (FWO: G065721N, G024925N). LVDB is supported by the Generet Award for Rare Diseases. PM has been awarded a Postdoctoral Mandate (PDM) from KU Leuven Internal Funds, is a recipient of the Early-Stage ALS Clinicians Grant from Target ALS, and has also received the Innovative Collaborative Projects with Biomarker Consortia Grant from Target ALS.

We thank the Nucleomics Core of VIB, KU Leuven (https://nucleomicscore.sites.vib.be/en) for the tissue procurement and sequencing.

## AUTHOR CONTRIBUTIONS

Conceptualization: P.V.D., D.R.T., A.S., P.M., M.G., Methodology: M.G., P.M., A.S., and P.V.D., Investigation: M.G., P.M., D.R.T., A.S and P.V.D., Data Curation: M.G and A.S., Formal analysis: M.G and A.S., Resources: D.R.T., K.P., and P.V.D., Validation: M.G., F.H., J.D., K.P., and P.V.D., Funding acquisition: P.V.D., Project administration: P.V.D., Supervision: A.S. and P.V.D., Visualization: M.G., Writing – original draft: M.G., A.S., and P.V.D., Writing – review & editing: All authors.

All corresponding authors have read and revised the manuscript, and fully agree with the accuracy of the results and its publication.

## DECLARATION OF GENERATIVE AI AND AI-ASSISTED TECHNOLOGIES

During the preparation of this work, the author(s) used AI tools, including ChatGPT (by OpenAI) and Gemini (by Google), to enhance the readability and language of the manuscript. No patient personal data or unpublished research findings were shared with these tools. The author(s) thoroughly reviewed and edited all AI-assisted content and take full responsibility for the final publication.

---

**Figure S1.**
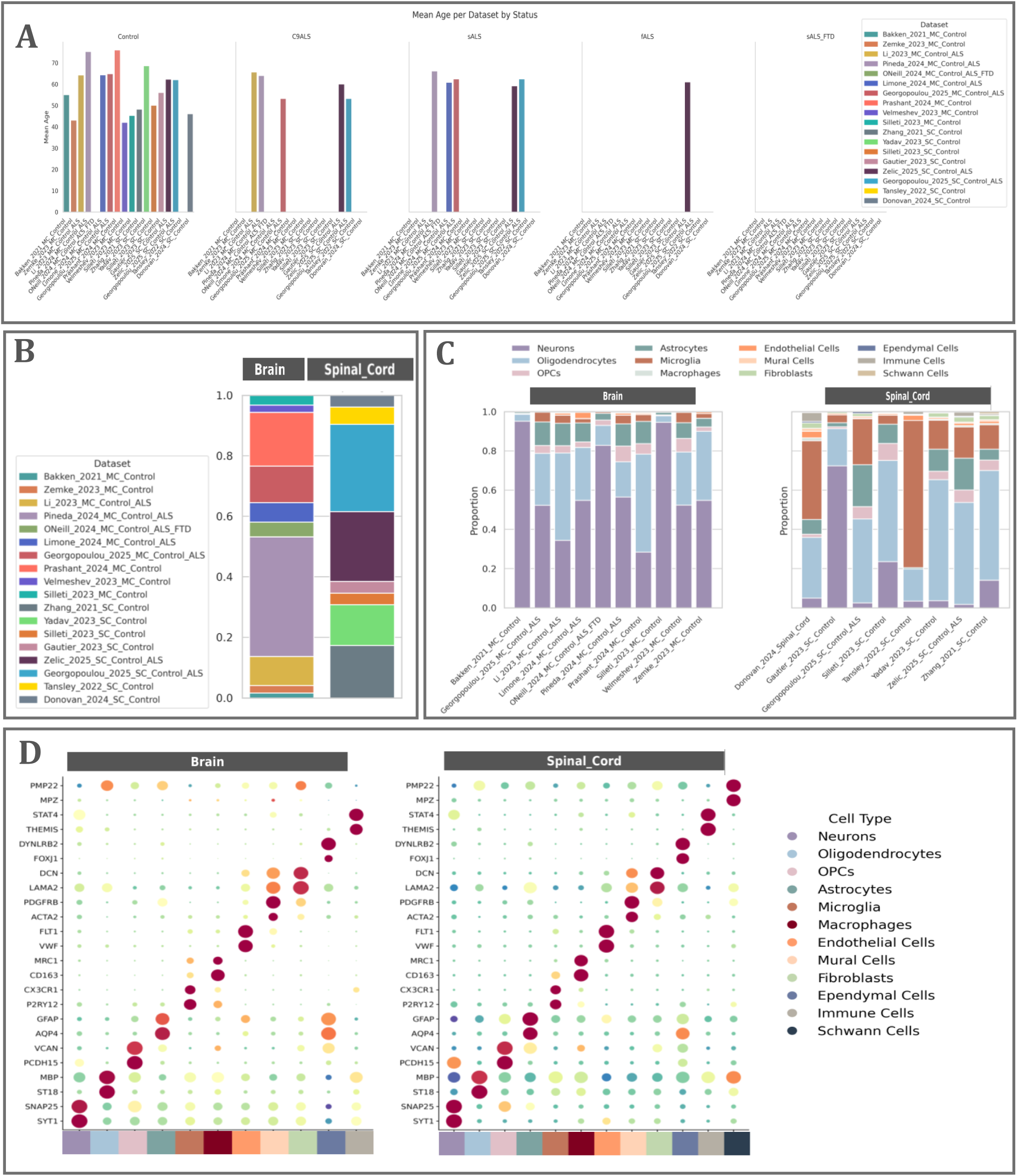
Clinical Data, Cell and Dataset Composition of the Motor Cortex and Spinal Cord Transcriptomic Atlas, Related to Figure 1. **(A)** Barplot of the mean age per dataset and per condition. **(B)** Barplot of the dataset composition in the motor cortex and the spinal cord. **(C)** Barplot of the cell type composition in the motor cortex and the spinal cord. Note that the Bakken et. al. and Gautier et. al. datasets are FACS-enriched for neurons, while the Tansley et. al. and Donovan et. al. datasets are FACS-enriched for microglia. **(D)** Dotplot of the main marker genes for each cell type in the motor cortex and the spinal cord.

**Figure S2.**
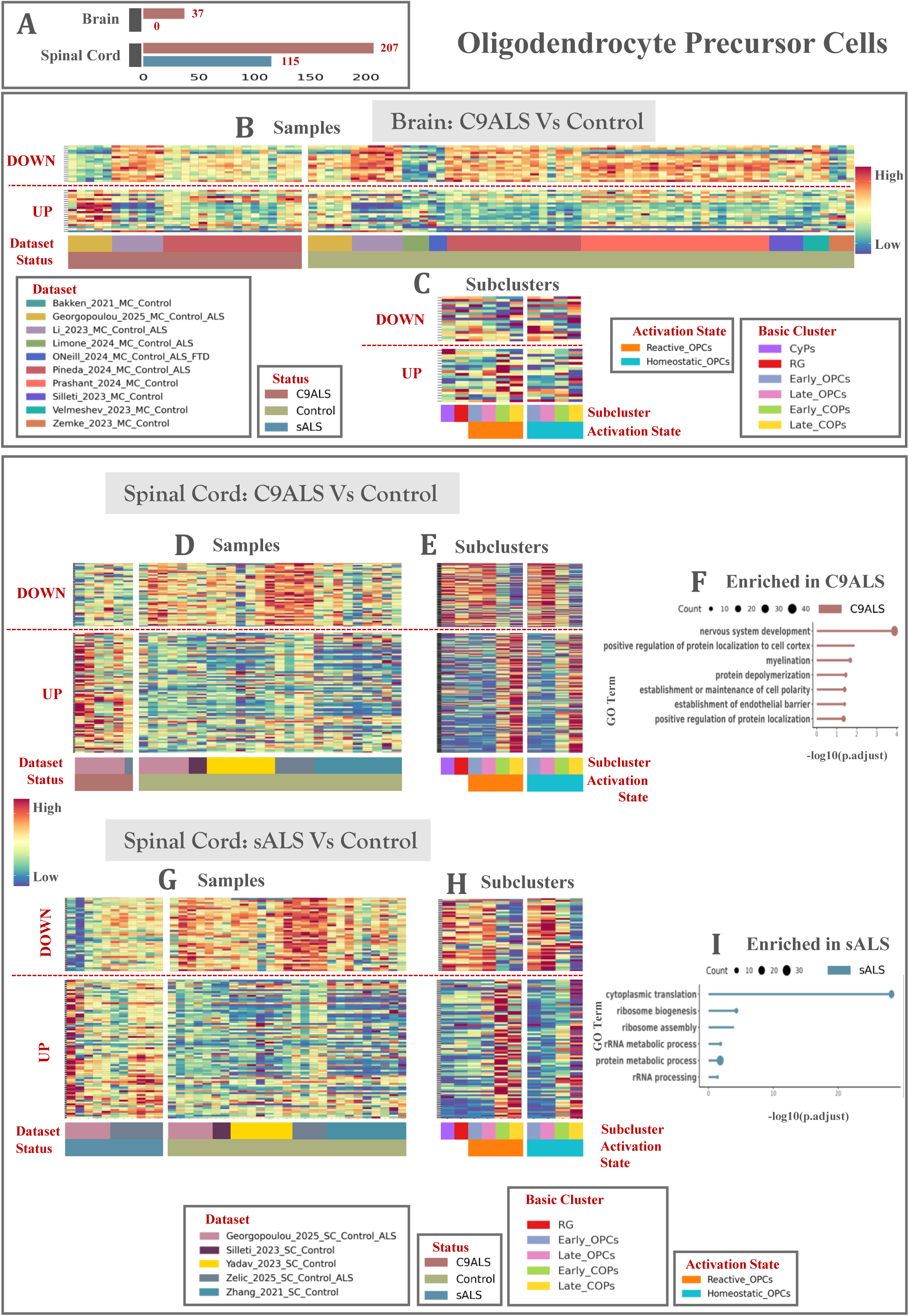
OPC Transcriptional Responses to ALS Are Region- and Condition-Dependent, Related to Figure 4. **(A)** Barplot of the amount of significant DEGs found in the motor cortex and the spinal cord with pseudobulk DESeq2 analysis and p.adj < 0.05. **(B)** MatrixPlot of the top up- and down-regulated transcripts in C9ALS vs. control in the motor cortex. Each column is a sample grouped by dataset and status. **(C)** MatrixPlot of the top up- and down-regulated transcripts in C9ALS vs. control in the motor cortex. Each column is a subcluster grouped by activation state. **(D)** MatrixPlot of the top up- and down-regulated transcripts in C9ALS vs. control in the spinal cord. Each column is a sample grouped by dataset and status. **(E)** MatrixPlot of the top up- and down-regulated transcripts in C9ALS vs. control in the spinal cord. Each column is a subcluster grouped by activation state. **(F)** Lollipop Plots of enriched and suppressed GO terms in C9ALS vs. control spinal cord. **(G)** MatrixPlot of the top up- and down-regulated transcripts in sALS vs. control in the spinal cord. Each column is a sample grouped by dataset and status. **(H)** MatrixPlot of the top up- and down-regulated transcripts in sALS vs. control in the spinal cord. Each column is a subcluster grouped by activation state. **(I)** Lollipop Plots of enriched and suppressed GO terms in sALS vs. control spinal cord. Abbreviations in this figure include: p.adj; p adjusted value, DEGs; differentially expressed genes.

**Figure S3.**
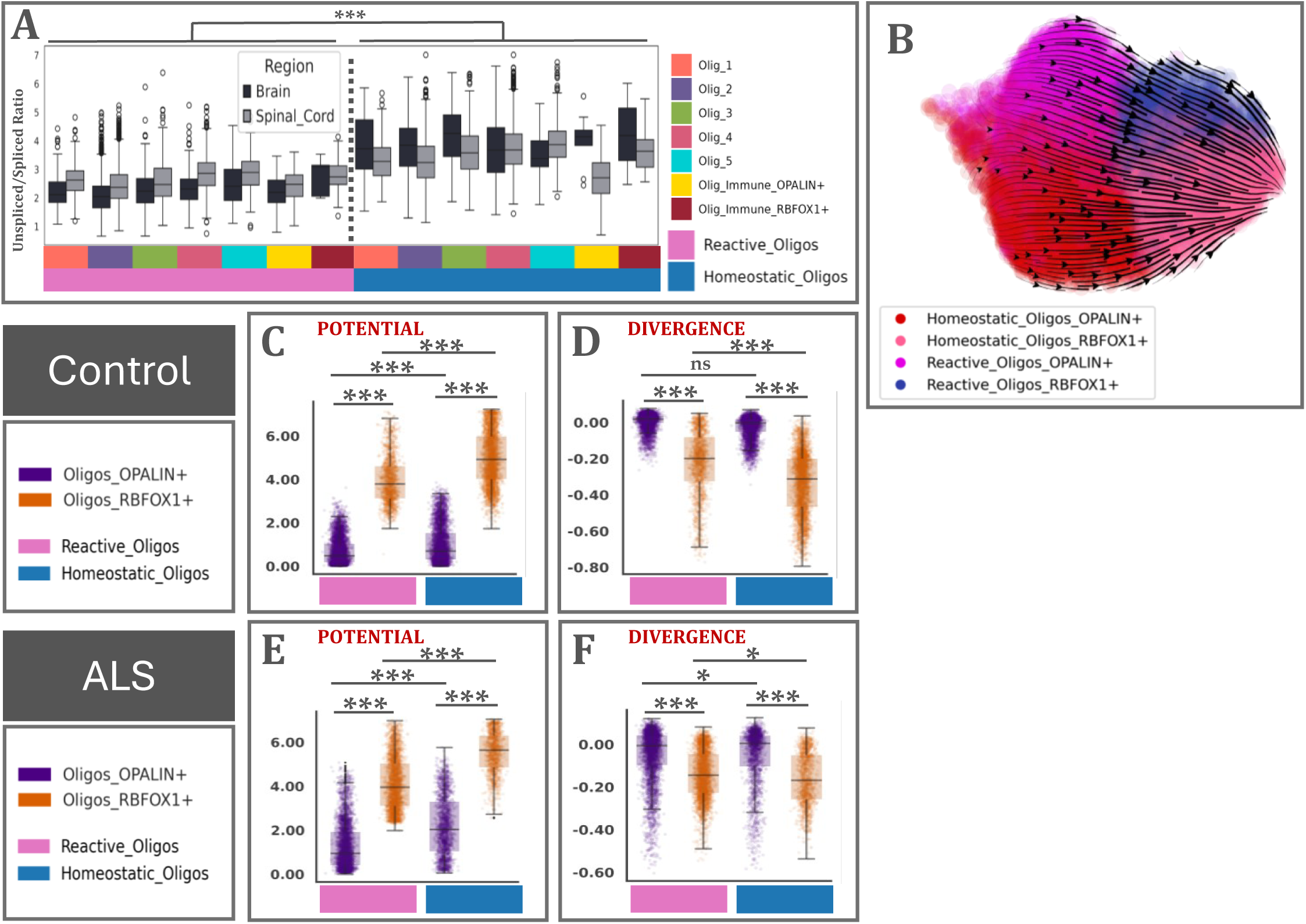
Splicing Kinetics and Vector Field Analysis Suggest a Potential Landscape of Oligodendrocyte States, Related to Figure 5. **(A)** Boxplot of the unspliced/spliced RNA ratio per subcluster grouped by activation state in the motor cortex and spinal cord control population. **(B)** Streamline plot of the predicted RNA velocity projected onto the UMAP embedding of the motor cortex and spinal cord control population. **(C)** Boxplot of the single-cell potential in oligodendrocytes from motor cortex and spinal cord control population, quantified using Velocyto. **(D)** Boxplot of the divergence in oligodendrocytes from control motor cortex and spinal cord control population, quantified using Velocyto. **(E)** Boxplot of the single-cell potential in oligodendrocytes from ALS (C9ALS and sALS) spinal cord, quantified using Velocyto. **(F)** Boxplot of the divergence in oligodendrocytes from ALS (C9ALS and sALS) spinal cord, quantified using Velocyto. llm; * p-value < 0.05, ** p-value < 0.01, *** p-value < 0.001

